# Predicting speech from a cortical hierarchy of event-based timescales

**DOI:** 10.1101/2020.12.19.423616

**Authors:** Lea-Maria Schmitt, Julia Erb, Sarah Tune, Anna Rysop, Gesa Hartwigsen, Jonas Obleser

**Affiliations:** Department of Psychology, University of Lübeck, Ratzeburger Allee 160,23562 Lübeck, Germany; Center of Brain, Behavior and Metabolism, University of Lübeck, Ratzeburger Allee 160, 23562 Lübeck, Germany; Lise Meitner Research Group Cognition and Plasticity, Max Planck Institute for Human Cognitive and Brain Sciences, Stephanstraße 1 A, 04103 Leipzig, Germany

## Abstract

How can anticipatory neural processes structure the temporal unfolding of context in our natural environment? We here provide evidence for a neural coding scheme that sparsely updates contextual representations at the boundary of events and gives rise to a hierarchical, multi-layered organization of predictive language comprehension. Training artificial neural networks to predict the next word in a story at five stacked timescales and then using model-based functional MRI, we observe a sparse, event-based “surprisal hierarchy”. The hierarchy evolved along a temporo-parietal pathway, with model-based surprisal at longest timescales represented in inferior parietal regions. Along this hierarchy, surprisal at any given timescale gated bottom-up and top-down connectivity to neighbouring timescales. In contrast, surprisal derived from a continuously updated context influenced temporo-parietal activity only at short timescales. Representing context in the form of increasingly coarse events constitutes a network architecture for making predictions that is both computationally efficient and semantically rich.

## Introduction

While the past predicts the future, not all context the past provides is equally informative: it might be outdated, contradictory, or even irrelevant. Nevertheless, the brain as a “prediction machine”^1^ is seemingly equipped with a versatile repertoire of computations to overcome these contextual ambiguities. A prominent example is speech, where a slip of the tongue may render the most recent context uninformative, but we can still predict the next word from its remaining context. At much longer time scales, we can re-use context that suddenly proves informative, as a speaker returns to a topic discussed earlier.

Using natural language comprehension as a working model, we here ask: How does the brain dynamically organize, evaluate, and update these complex contextual dependencies over time to make accurate predictions?

A robust principle in cerebral cortex is the decomposition of temporal context into its constituent timescales along a hierarchy from lower to higher-order areas, which is evident across species^2,3^, recording modalities^4,5^, sensory modalities^6,7^, and cognitive functions^8,9^. For instance, sensory cortices closely track rapid fluctuations of stimulus features and operate on short timescales (e.g., Ref.^10^). By contrast, association cortices integrate stimuli over an extended period and operate on longer timescales (e.g., Ref.^11^).

Conceptually, such hierarchies of “temporal receptive windows” are often subsumed under the framework of predictive coding^12^: A nested set of timescale-specific generative models informs predictions on upcoming sensory input and is updated based on the actual input^13^. In particular, context is thought to shape the prediction of incoming stimuli via feedback connections. These connections would link each timescale to its immediate shorter timescale, while the prediction error is propagated forward through the hierarchy^1,14^. Indeed, hierarchical specialization has been shown empirically to emerge from structural and functional large-scale connectivity across cortex^15,16^. More precisely, feedforward and feedback connections^17,18^ shown to carry prediction errors and predictions^19,20^, respectively, are a hallmark of hierarchical predictive coding.

However, studies on the neural underpinnings of predictive coding have primarily used artificial stimuli of short temporal context (but see Ref. ^21^) and employed local vs. global violations of expectations, effectively manifesting a two-level cortical hierarchy (but see Ref.^22^). We thus lack understanding whether the hierarchical organization of prediction processes extends to natural environments unfolding their temporal dependencies over a multitude of interrelated timescales.

With respect to functional organization in cortex, temporo-parietal areas are sensitive to a rich set of hierarchies and timescales in speech^23-27^. Most relevant to the present work, semantic context in a spoken story has been shown to map onto a gradient extending from early auditory cortex representative of words up to intraparietal sulcus, representative of paragraphs^28^. This timescale-specific representation of context is reminiscent of the multi-layered generative models proposed to underlie predictive coding^29,30^. Compatible with this notion, previous studies on speech comprehension found evidence for neural coding of prediction errors at the level of syllables^31^, words^32^, or discourse^33^.

Yet the interactions between multiple representational levels of speech in predicting upcoming words remain unclear. Here, we ask whether the processing hierarchy enabling natural speech comprehension is also implicated in evaluating the predictiveness of timescale-specific semantic context and integrating informative context into predictions.

We do not know how context that unfolds at a particular timescale would be updated cortically when the listener receives new bottom-up input. One attractively simple candidate architecture is the *continuously updating processing hierarchy*. In a recent study, Chien and Honey^34^ showed that neural responses to a story rapidly aligned across participants in areas with shorter, but only later in areas with longer receptive windows. This response pattern was best explained by a computational model which immediately integrates upcoming input with context representations at all timescales. An important implication of such continuous updates is that all context representations are continuously tuned to current processing demands.

A competing candidate architecture, however, is the *sparsely updating processing hierarchy*. For example, it is known that scenes in a movie are encoded as event-specific neural responses^35^ and that more parietal receptive windows represent increasingly coarse events in movies^36^. Such an event hierarchy is effectively based on the boundaries of events: It calls for stable working memory representations that are *sparsely* recombined with preceding events at higher processing stages only at the end of an event. The simultaneous representation of distinct events in working memory allows to draw on diverse context when making predictions. We here hypothesize that such a sparsely updating network architecture is a more appropriate model for prediction processes in the brain.

In the present study, we recorded blood-oxygen-level-dependent (BOLD) responses while participants listened to a narrated story, which provides rich semantic context and captures the full dynamic range of speech predictability^37^. Following the rationale that neural computations can be inferred by comparing the fit of neural data to outputs from artificial neural networks with different architectures^38,39^, we derived context-specific surprisal associated with each word in the story from single layers of long short-term memory (LSTM)-based language models with either a continuous or a sparse updating rule.

Here, we show that the event-based organization of semantic context provides a valid model of predictive processing in the brain. We show that a “surprisal hierarchy” of increasingly coarse event timescales evolves along the temporo-parietal pathway, with stronger connectivity to neighbouring timescales in states of higher word surprisal. Surprisal derived from continuously updated context had a (non-hierarchical) effect on temporo-parietal activity only at short timescales. Our results suggest that representing context in the form of increasingly coarse events constitutes a network architecture that is both, computationally efficient and semantically rich for making predictions.

## Results

Thirty-four participants listened to a one-hour narrated story while their hemodynamic brain responses were recorded using functional magnetic resonance imaging (fMRI). To emulate a challenging listening scenario, the story audio was presented against a competing stream of resynthesized natural sounds (for an analysis focussing on this acoustic representation see Ref.^40^).

The surprisal associated with each word in the story was modelled at multiple timescales of semantic context by two artificial neural networks, one with a continuous updating rule (LSTM) and another one with a sparse updating rule (HM-LSTM; Figure 1A).

**Figure 1.**
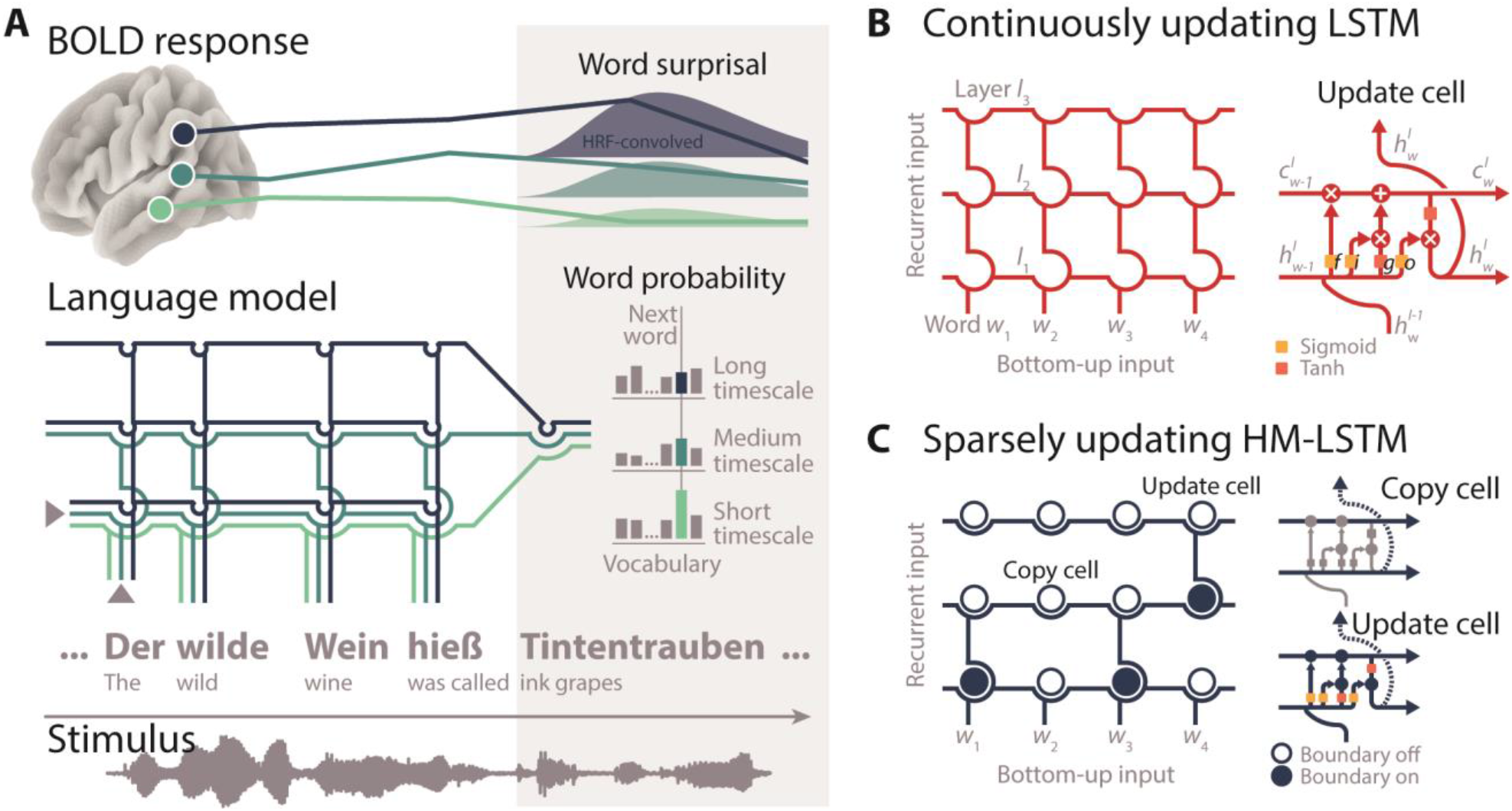
Modelling neural speech prediction with artificial neural networks. **(A)** Participants listened to a story (grey waveform) during fMRI. Based on its semantic context (“The wild wine was called”), a language model predicted each word in the story (“ink grapes”). The probability of the next word was read out from each layer of the model separately, with higher layers accumulating information across longer semantic timescales. Word probabilities were transformed to surprisal, convolved with the hemodynamic response function and mapped onto temporo-parietal BOLD time series. **(B)** Two language models were trained. With each new word-level input, the “continuously updating” long short-term memory (LSTM)^41^ combines “old” recurrent long-term 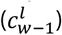 and short-term memory states 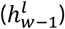 with “new” bottom-up semantic input 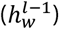 at each layer *I*. This allowed semantic information to continuously flow to higher layers with each incoming word, f: forget gate, i: input gate, g: candidate state, o: output gate. **(C)** The “sparsely updating” hierarchical multiscale LSTM (HM-LSTM)^42^ was designed to learn the hierarchical structure of text. An upper layer keeps its representation of context unchanged (copy mechanism) until a boundary indicates the end of a timescale on the lower layer and information is passed to the upper layer (update mechanism). Networks were unrolled for illustration only.

First, we encoded surprisal at multiple timescales into univariate neural responses and fit a gradient to temporo-parietal peak locations of timescales. Second, we decoded timescale surprisal from patterns of neural responses in single parcels and compared decoding accuracies between language models. Finally, we investigated how surprisal gates the information flow between brain regions sensitive to different timescales.

All encoding and decoding models were estimated separately per each language model, using ridge regression with four-fold cross-validation.

### Two competing language models of hierarchical speech prediction

We trained two artificial neural networks on more than 130 million words of running text to predict an upcoming word by its preceding semantic context. More specifically, language models consisted of long short-term memory cells (LSTM)^41^, which incorporate context that might become relevant at some time (*cell state*) or that is relevant already to the prediction of the next word (*hidden state*). By stacking five LSTM layers, our models operated on different timescales of context, with higher layers coding for long-term dependencies between words.

In the continuously updating (or “vanilla”) LSTM, recurrent memory states are updated at each layer with every new bottom-up word input (Figure 1B). A second model, the *hierarchical multiscale* LSTM^42^, referred to as “sparsely updating HM-LSTM”, employs a revised updating rule where information from a lower layer is only fed forward at the end of its representing timescale (Figure 1C). This allows for less frequent updates between layers and stronger separation between contextual information represented at different layers.

### Three model-derived metrics of predictiveness at multiple timescales

For each word in the entire presented story (> 9,000 words), we determined its predictability given the semantic context of the preceding 500 words. Hidden states were combined across layers and mapped to an output module, which denotes the probability of occurring next for every word in a large vocabulary of candidate words. The word with the highest probability was selected as the *predicted* next word. Overall, the LSTM (proportion correct across words: 0.13) and the HM-LSTM (0.12; Supplementary Figure 1) were on par in accurately predicting the next word in the story.

To derive the predictability of words based on layer-specific context (or, for our purpose, timescale), we allowed information to freely flow through pre-trained networks, yet only mapped the hidden state of one layer to the output module by setting all other network weights to zero. Outputs from these “lesioned” language models represented the five timescales.

As the primary metric of predictiveness, we calculated the degree of surprisal associated with the occurrence of a word given its context (i.e., negative logarithm of the probability assigned to the *actual* next word). The surprisal evoked by an incoming word indexes the amount of information that was not predictable from the context represented at a specific timescale^43,44^. Of note, surprisal was considerably higher for longer timescales in the LSTM (*p* < 0.001, Cohen’s *d* = 2.43; compared to slopes drawn from surprisal shuffled across timescales) but remained stable across timescales in the HM-LSTM (*p* = 0.955, *d* = 0.05; direct comparison LSTM vs. HM-LSTM: *p* < 0.001, *d* = 2.7; Figure 2).

**Figure 2.**
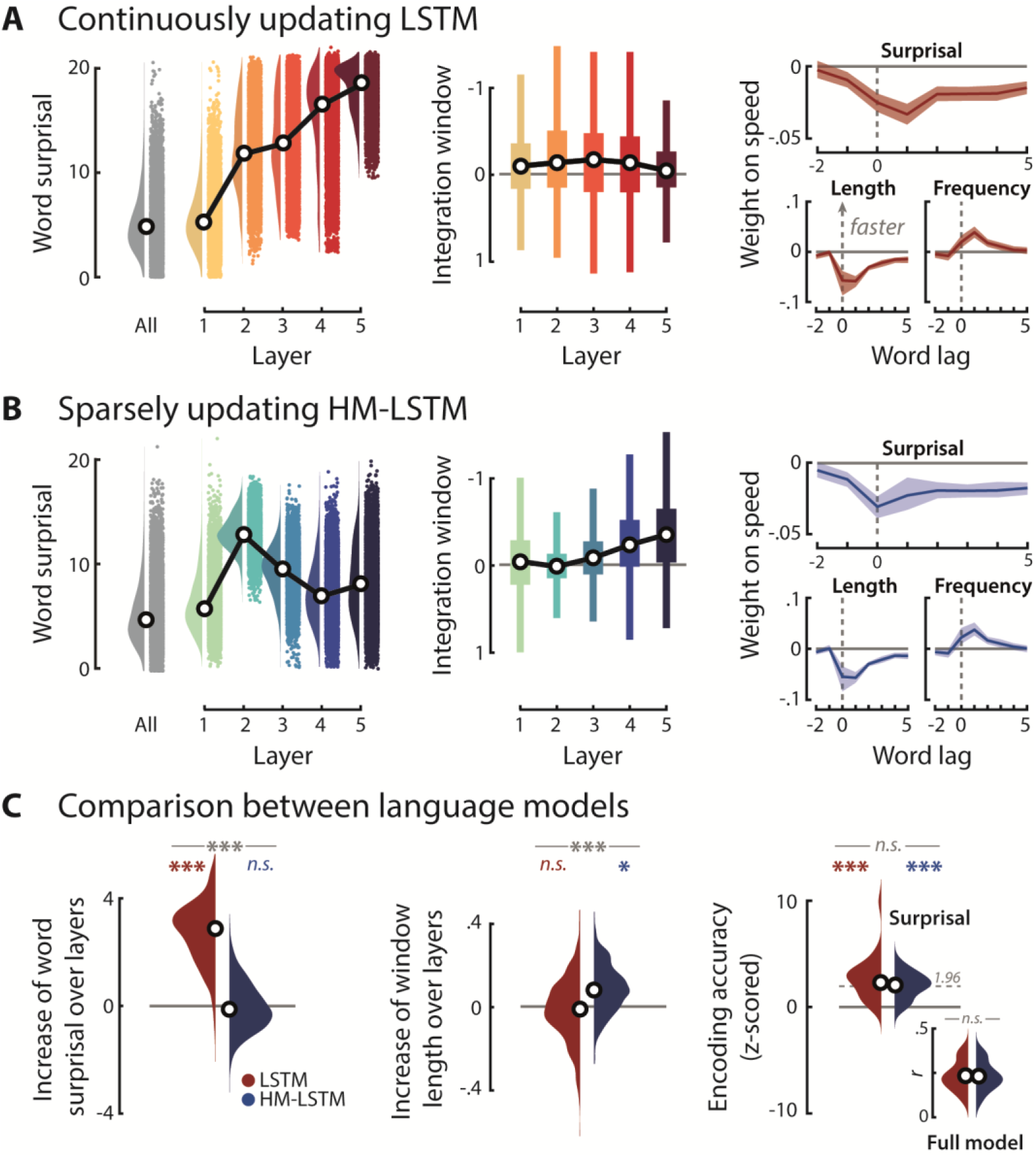
Evaluating model-derived surprisal. **(A)** Left: Word surprisal derived from the “full” LSTM model including all layers (grey distribution) and from single layers of “lesioned” LSTM models (coloured distributions); black circles represent grand-median surprisal. Middle: Input to the LSTM was scrambled at different granularities corresponding to an increase in the length of intact context (i.e., 1–256 words). For each layer of the LSTM, linear functions were fit to word surprisal across these context windows. A negative slope parameter indicates a stronger benefit (or lower surprisal) from longer context (i.e., larger integration window). Right: Speed in a self-paced reading task was modelled as a function of time-lagged predictiveness and a set of nuisance regressors. Weight profiles illustrate the temporal dynamics of the surprisal effect in the full model (top) in comparison to word length (bottom, left) and word frequency (bottom, right); positive weights indicate an increase in response speed; error bands represent ±SEM. **(B)** Same as in A, but for the sparsely updating HM-LSTM. **(C)** Left: We fit linear functions to word surprisal across layers and compared resulting slope parameters to null distributions drawn from shuffled layers and between LSTM (red) and HM-LSTM (blue). Middle: Linear fit to integration windows across layers, indicating the benefit of higher layers from longer context. Right: Encoding accuracy in the self-paced reading task uniquely explained by the predictiveness of context (standardized to scrambled features of predictiveness); dotted grey line indicates critical significance level for single participants. Inset shows non-standardized encoding accuracies. *** *p* < 0.001, * *p* < 0.05, n.s.: not significant.

To determine the temporal integration window of each timescale, we scrambled input to the network at different granularities corresponding to a binary logarithmic increase in the length of intact context (i.e., 1-256 words). The LSTM showed no such increase of temporal integration windows at higher layers (LSTM: *p* = 0.219, *d* = 0.11). In contrast, in the HM-LSTM, surprisal decreased more strongly at longer compared to shorter timescales as more intact context became available (HM-LSTM: *p* = 0.027, *d* = 0.73; LSTM vs. HM-LSTM: *p* < 0.001, *d* = 0.76; Figure 2).

Our secondary metrics expressed the predictability of a word in relation to other words, that is, (1) the entropy of the probability distribution predicted for individual words (indicative of the difficulty to make a definite prediction) and (2) the dissimilarity of vector representations (or embeddings) coding for the constituent linguistic features of the predicted and actual next word (Product-moment correlation; indicative of conceptual (un-)relatedness).

We derived surprisal, entropy and dissimilarity associated with single words from “lesioned” models at each of five timescale and from “full” models across all timescales, separately for each language model. All features were convolved with the hemodynamic response function, and we will collectively refer to them as “features of predictiveness” from here on.

### Higher model-derived surprisal of words slows down reading

To test the behavioural relevance of model-based predictiveness, another 26 participants performed a self-paced reading task where they read the transcribed story word-by-word on a noncumulative display and pressed a button as soon as they had finished reading.

When regressing response speed onto time-lagged features of predictiveness and a set of nuisance regressors (e.g., word length and frequency), we found that—as expected—reading speed slowed down for words determined as more surprising by language models given the full context across all timescales (Figure 2A).

Further, we predicted response speed on held-out testing data and z-scored the resulting encoding accuracy (i.e., Product-moment correlation of predicted and actual response speed) to a null distribution drawn from scrambled features of predictiveness while only keeping nuisance regressors intact. This yielded the unique contribution of the predictiveness of words (i.e., surprisal, entropy and dissimilarity) to reading speed, which was significant for both language models (LSTM: *p* < 0.001, *d* = 1.51; HM-LSTM: *p* < 0.001, *d* = 1.64, LSTM vs. HM-LSTM: *p* = 0.975, *d* = 0.35). Together, these findings suggest that both language models picked up on processes of speech prediction that shape behaviour.

### Selecting temporo-parietal regions of interest involved in speech processing

We hypothesized that the speech prediction hierarchy is represented as a gradient along the temporo-parietal pathway. This rather coarse region of interest was further refined to only include regions implicated in processing of the listening task.

To this aim, we calculated pairwise intersubject correlations^45^, which revealed consistent cortical activity across participants in a broad bilateral language network. Responses were most prominently shared in auditory association cortex and lateral temporal cortex as well as premotor cortex, paracentral lobule and mid cingulate cortex (Figure 3A). Crucially, as sound textures presented in the competing stream were randomly ordered across participants, this approach allowed us to extract shared responses specific to the speech stream.

**Figure 3.**
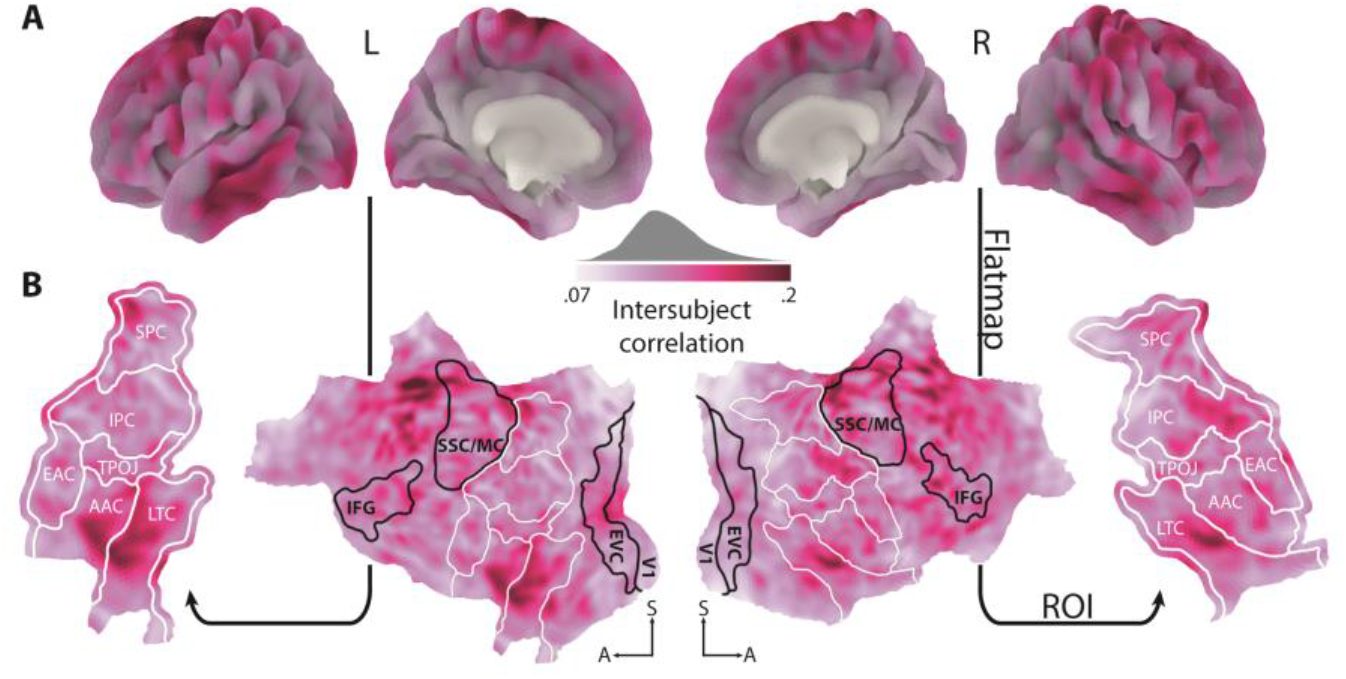
Selection of regions of interest. **(A)** When listening to a story against background noise, pairwise intersubject correlations showed stronger synchronization of BOLD activity in cortical areas implicated in the language network. **(B)** The cortical surface was flattened. All temporal and parietal parcels^46^ highlighted by white outlines were included as regions of interest (ROI) in the following analyses. Black outlined parcels serve as reference point, only. EAC: early auditory cortex, AAC: auditory association cortex, LTC: lateral temporal cortex, TPOJ: temporo-parieto-occipital junction, IPC: inferior parietal cortex, SPC: superior parietal cortex, V1: primary auditory cortex, EVC: early visual cortex, IFG: inferior frontal gyrus, SSC/MC: somatosensory and motor cortex. Maps were smoothed with an 8 mm FWHM Gaussian kernel for illustration only.

All further analyses were limited to those parcels in temporal and parietal cortex^46^ that showed significant median intersubject correlations in more than 80% of vertices (*p_FDR_* < 0.01; ranked against a bootstrapped null distribution). The cortical sheet of the six parcels determined as regions of interest (ROI) was flattened, resulting in a two-dimensional plane spanned by an anterior-posterior and inferior-superior axis (Figure 3B).

We expected gradients of speech prediction to unfold along the inferior-superior axis, that is, from temporal to parietal areas.

### Differential tuning to continuously and sparsely updated timescales of surprisal in temporoparietal cortex

After the predictiveness of speech was encoded into neural activity of single vertices, we extracted temporo-parietal weight maps of word surprisal at each timescale for both language models.

When performing spatial clustering on these weight maps (*p*_vertex_ and *p*_cluster_ < 0.05; compared to scrambled surprisal by means of a cluster-based permutation test), we found large positive clusters in both hemispheres for shorter timescales of the LSTM (Figure 4A, yellow outlines) but, if at all, only focal clusters for longer timescales (Figure 4A, red outlines). Hence, temporo-parietal activity primarily increased in response to words that were less predictable by the context provided at shorter, continuously updated timescales. In contrast, clusters of distinct polarity, location and extent were observed for all timescales of the HM-LSTM (Figure 4B), suggesting that even longer timescales had the potency to modulate temporo-parietal activity when they were sparsely updated.

**Figure 4.**
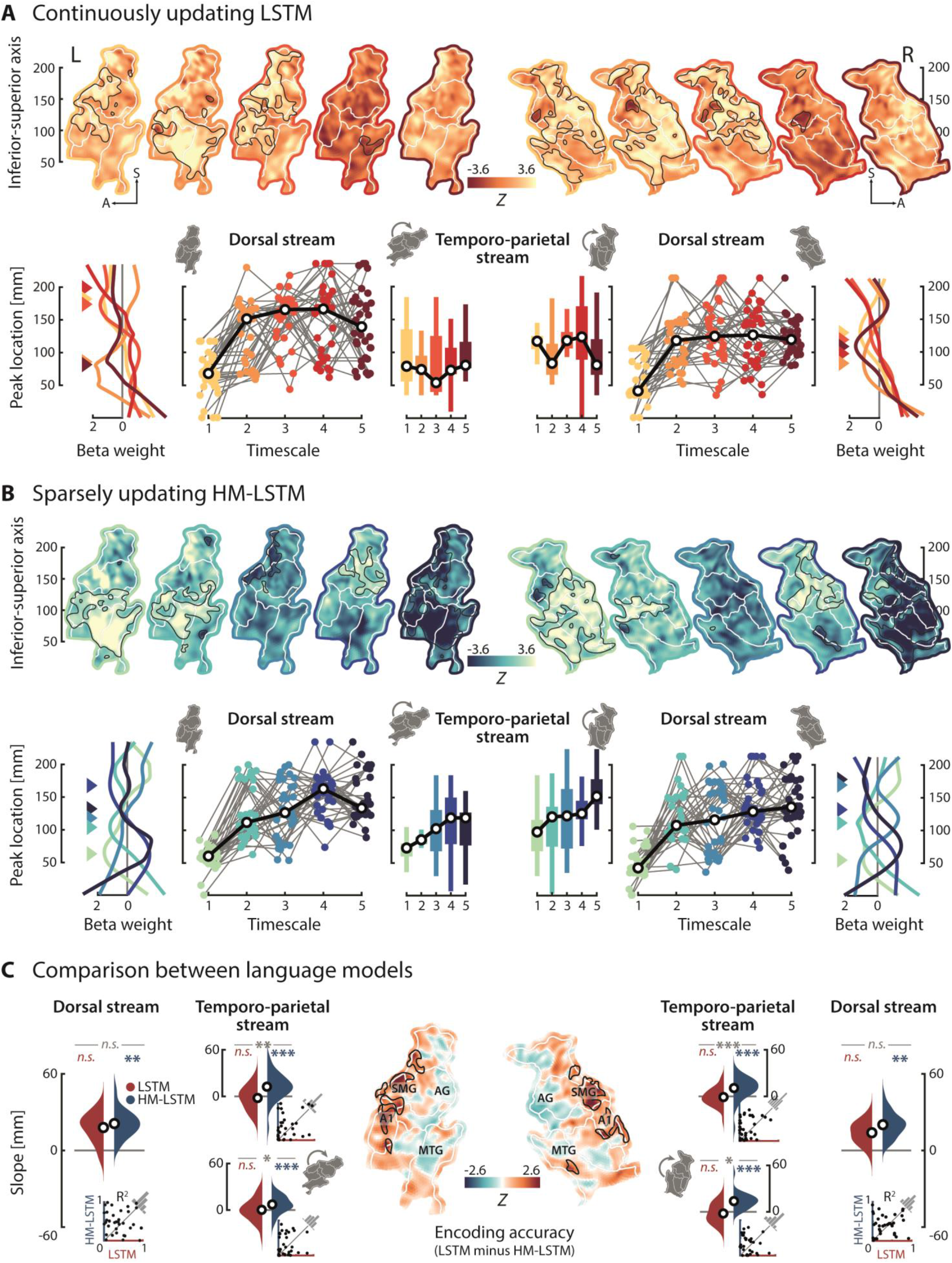
Encoding the timescales of surprisal. **(A)** Temporo-parietal BOLD time series were mapped onto the predictiveness of speech derived at five timescales from the continuously updating LSTM. Top row: Maps show z-values from timescale-specific weights of surprisal tested against a null distribution drawn from scrambled surprisal; black outlines indicate significant clusters; white outlines indicate parcels; coloured outlines indicate short (light yellow) to long (dark red) timescales, separately for the left and right hemisphere. Bottom row: For each timescale, we determined its peak coordinate along the inferior-superior axis (coloured triangles), here shown for grand-average weight profiles for illustration only. Testing for a processing hierarchy along the dorsal stream, timescales were restricted to peak superior to the first timescale; coloured dots connected by grey lines represent peak coordinates of single participants; black circles represent grand-median peak coordinates. Testing for a processing hierarchy along the temporo-parietal stream, timescales were allowed to peak at any location. Maps were rotated by −45° to test simultaneous effects on the inferior-superior and anterior-posterior axis in the left hemisphere. Note that right-hemispheric maps already had an initial rotation off the inferior-superior axis, so rotating these maps resulted in testing for effects on the inferior-superior axis only. **(B)** Encoding maps and timescale-specific peak locations for the sparsely updating HM-LSTM. **(C)** Linear functions were fit to peak coordinates across timescales and resulting slope parameters were compared to null distributions drawn from scrambled coordinates and between language models, separately for the dorsal (not rotated) and temporo-parietal stream (top: not rotated; bottom: rotated) in both hemispheres. Black circles represent grand-average slope parameters; insets depict coefficients of determination for linear fits of single participants. Temporo-parietal encoding accuracies displayed on ROI maps were z-scored to null distributions drawn from scrambled features of predictiveness and compared between language models; black outlines indicate significant clusters; maps were smoothed with an 8 mm FWHM Gaussian kernel for illustration only; SMG: supramarginal gyrus, AG: angular gyrus, A1: primary auditory cortex; MTG: middle temporal gyrus, * *p* < 0.05, ** *p* < 0.01, *** *p* < 0.001, *n.s*.: not significant.

### Sparsely updated timescales of surprisal evolve along a temporo-parietal processing hierarchy

To probe the organization of timescales along a temporo-parietal gradient, we collapsed across the anterior-posterior axis of weight maps and selected the local maximum with the largest positive value on the inferior-superior axis. Fitting a linear function to those peak coordinates of timescales, we found flat slope parameters indicating random ordering of LSTM timescales in both hemispheres (left: *p* = 0.458, *d* = −0.15; right: *p* = 0.716, *d* = −0.07; compared to slopes drawn from coordinates scrambled across timescales, Figure 4C). Conversely, we found steep positive slopes for the HM-LSTM in both hemispheres (left: *p* < 0.001, *d* = 0.72; right: *p* < 0.001, *d* = 0.75; Figure 4C), reflecting the representation of longer timescales in more parietal regions. On average, the left hemisphere represented sparsely updated timescales 12 mm superior (along the unfolded cortical surface) to their directly preceding timescale. Most relevant, this finding was underpinned by a significant difference of slope parameters between the LSTM and HM-LSTM (left: *p* = 0.005, *d* = 0.9; right: *p* < 0.001, *d* = 0.89; Figure 4C), demonstrating a temporo-parietal processing hierarchy of word surprisal that preferably operates on sparsely updated timescales.

The absence of a gradient for continuously updated timescales was corroborated when specifically targeting the dorsal processing stream. To this aim, we confined the first timescale to peak in temporal regions and all other timescales to peak superior to the first timescale in more parietal regions. Slope effects along the dorsal stream largely complied with those found along the inferior-superior axis (LSTM: all *p* ≥ 0.134; HM-LSTM: all *p* ≤ 0.006; LSTM vs. HM-LSTM: all *p* ≥ 0.103), thereby ruling out the possibility that the presence of a competing ventral stream obscured the consistent ordering of timescales.

Additionally, rotating weight maps by −45° before collapsing across the first dimension showed that sparsely updated timescales were not only processed along an inferior-superior but also an anterior-posterior gradient in the left hemisphere (LSTM: *p* = 0.921, *d* = 0.02; HM-LSTM: *p* = 0.001, *d* = 0.65; LSTM vs. HM-LSTM: *p* = 0.011, *d* = 0.55). As right-hemispheric maps already had a strong initial rotation off the inferior-superior axis, rotating these maps merely confirmed that longer timescales are processed in more superior regions (LSTM: *p* = 0.355, *d* = −0.18; HM-LSTM: *p* < 0.001, *d* = 1.39; LSTM vs. HM-LSTM: *p* = 0.039, *d* = 1.45).

Unlike for the sparsely updated timescales of surprisal, neither the timescales of entropy (all *p* ≥ 0.583) nor dissimilarity (all *p* ≥ 0.623) organized along a dorsal gradient (Supplementary Figure 2). Further, effects of HM-LSTM timescale surprisal were dissociable from a simple measure of semantic incongruence between words in the story and their context at five timescales logarithmically increasing in length (all *p* ≥ 0.5; Product-moment correlation of target and average context embedding). This highlights the specificity of the observed gradient to prediction processes in general and word surprisal in particular.

### A segregated stream of continuously updated timescales of surprisal?

To determine the contribution of predictiveness to overall encoding accuracy on held-out data, we z-scored accuracies relative to null distributions drawn from scrambled features of predictiveness while keeping additional (spectro-temporal) acoustic and linguistic nuisance regressors intact. Interestingly, the LSTM produced—in comparison to the HM-LSTM—better predictions in early auditory cortex and supramarginal gyrus (*p*_vertex_ and *p*_cluster_ < 0.05; cluster-based permutation paired-sample *t*-test; Figure 4C). On the other hand, predictions of the HM-LSTM seemed slightly more accurate than for the LSTM along middle temporal gyrus, temporo-parieto-occipital junction and angular gyrus, even though not statistically significant. Taking into account the broad clusters found earlier specifically for shorter (but not longer) LSTM timescales, this poses the question whether continuously updated timescales take full effect only in earlier medial temporal and anterior parietal processing stages, whereas the sparsely updating processing hierarchy evolves along a separate lateral temporal and posterior parietal route.

### Parietal regions preferentially represent sparsely updated, long timescales

In a complementary decoding approach, we reconstructed the timescales of surprisal from patterns of neural activity in single regions of interest (i.e., temporo-parietal parcels). Reconstructed timescales were z-scored to scrambled features of predictiveness. Overall, bilateral auditory association cortex and lateral temporal cortex yielded highest decoding accuracies on held-out testing data for both language models (Figure 5A). More parietal regions showed comparably lower decoding accuracies. Nevertheless, these accuracies can be deemed meaningful, as the pattern mirrors the lower intersubject correlations in parietal compared to temporal regions (Figure 3), which are commonly found during natural listening (e.g., Ref. ^47-49^). This indicates an overall greater variability of neural responses in parietal regions irrespective of the timescales of surprisal.

**Figure 5.**
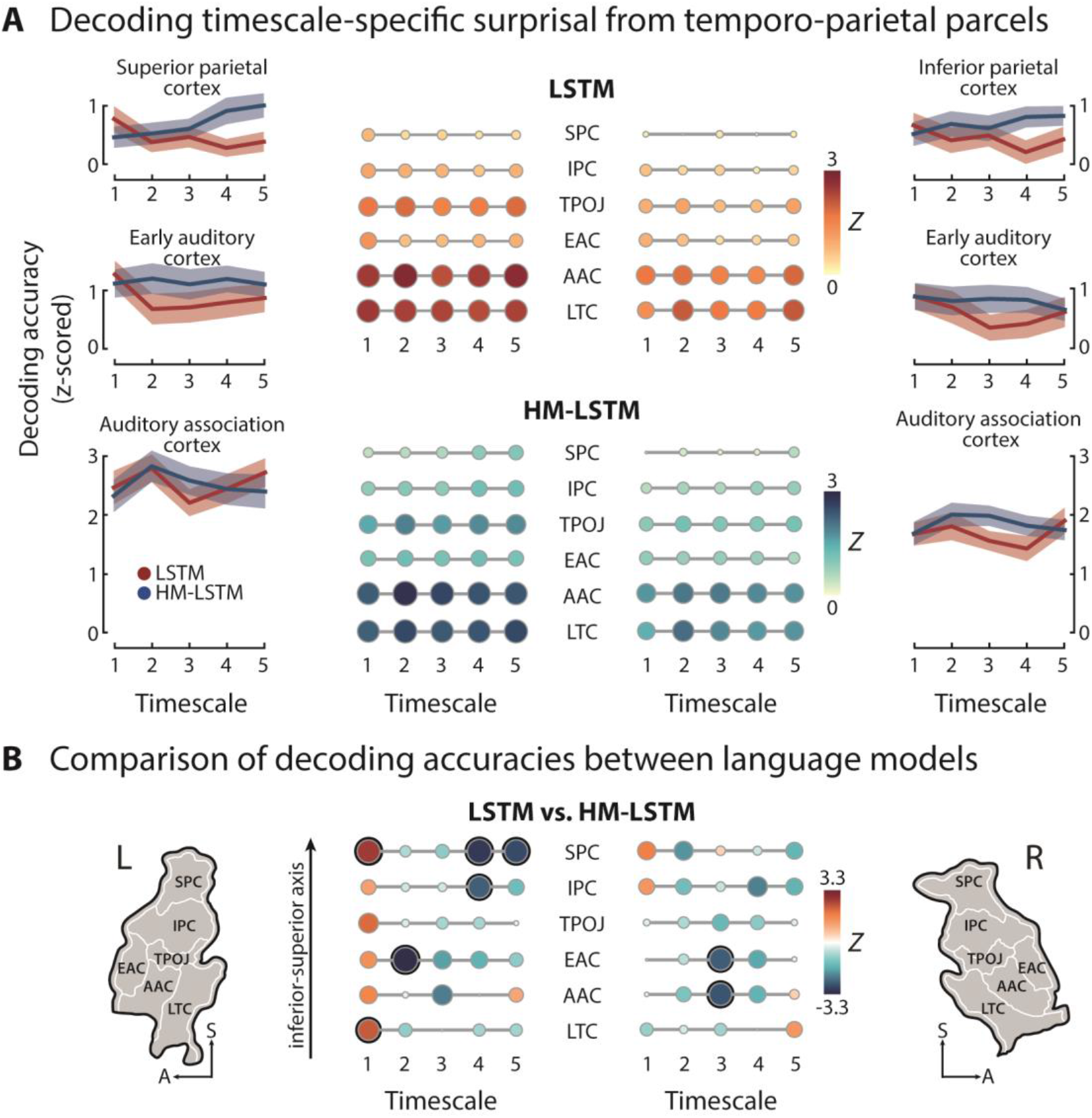
Decoding the timescales of surprisal. **(A)** Surprisal at different timescales was decoded from regions of interest. Matrices depict decoding accuracies determined on held-out testing data and z-scored to null distributions drawn from scrambled surprisal, separately for the LSTM (top) and HM-LSTM (bottom) in the left and right hemisphere; colour and size of circles scale to decoding accuracy. Of note, some z-scored decoding accuracies in more superior parcels fell below an average value of 1.96. However, z-scores were indicative of significance only on the level of single participants. Line plots illustrate patterns of decoding accuracies across timescales in select regions of interest; error bands represent ±SEM. **(B)** Decoding accuracies were contrasted between language models by means of a permutation test on the mean of differences; black circles indicate *P*_FDR_ < 0.05; maps indicate location of parcels. EAC: early auditory cortex, AAC: auditory association cortex, LTC: lateral temporal cortex, TPOJ: temporo-parieto-occipital junction, IPC: inferior parietal cortex, SPC: superior parietal cortex.

Contrasting decoding accuracies between language models, temporo-parietal regions of interest showed an overall preference for the shortest LSTM but longer HM-LSTM timescales (Figure 5B). This suggests that the predominance of the LSTM in early auditory cortex and supramarginal gyrus observed for encoding accuracies of the encoding model is specific to the shortest timescale, while lateral temporal and posterior parietal regions reflect longer timescales of the HM-LSTM, lending further support to the functional dissociation of two routes of predictive processing in speech prediction.

In particular, left-hemispheric early auditory cortex contained more information on medium, sparsely updated than continuously updated timescales, whereas inferior and superior parietal cortex preferentially represented long, sparsely updated timescales (*p*_FDR_ < 0.05). This finding converges with the organization of the gradient described earlier for sparsely updated but not continuously updated timescales.

### Surprisal at sparsely updated timescales gates connectivity along the processing hierarchy

After establishing the temporo-parietal processing hierarchy, we examined the modulatory effect of surprisal on connectivity between peak locations of timescales taken from the encoding analysis. To this aim, we created psychophysiological interactions between the BOLD response at the peak location of one timescale and word surprisal at the same timescale. The BOLD response of each (target) timescale was mapped onto psychophysiological interactions of all other (predictor) timescales (Figure 6A).

**Figure 6.**
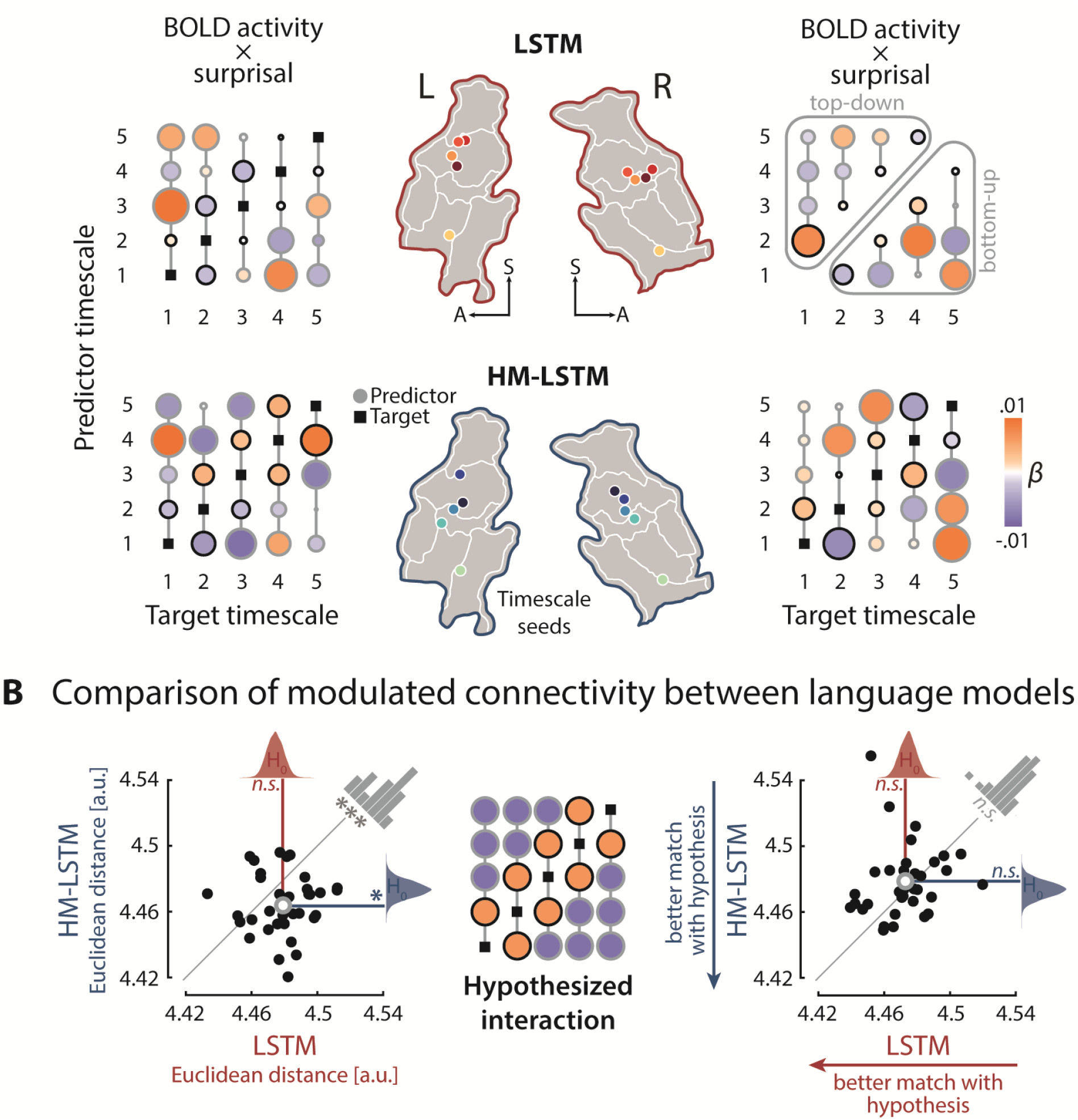
Surprisal-dependent modulation of effective connectivity. **(A)** A sphere of 6 mm was centred on median peak locations of timescales as defined in the encoding analysis (coloured circles on temporo-parietal maps) and BOLD responses were averaged within these timescale seeds. BOLD time series at one (target) timescale were regressed onto psychophysiological interactions of all other (predictor) timescales (i.e., pointwise product of timescale-specific BOLD and surprisal time series). For each target seed, we added a column vector of timescale-specific predictor weights to a 5-by-5 matrix with an empty main diagonal. Matrices were created separately for each language model (top: LSTM, bottom: HM-LSTM) and hemisphere. The upper triangle of a matrix indicates top-down, the lower triangle bottom-up information flow; black outlines of circles indicate timescale pairing for which we expected an increase of connectivity, a decrease of connectivity was expected for pairings with grey outlines (see hypothesized interaction pattern in 6B). **(B)** A hypothesized matrix of psychophysiological interactions was created, with positive weights on diagonals below and above the main diagonal, indicating increased connectivity between neighbouring timescales when surprisal is high. The Euclidean distance between observed and hypothesized matrices was compared to null distributions of distances drawn from target timescales shifted in time (coloured density plots), separately for the left and right hemisphere; black dots indicate distances of single participants; grey circles indicate mean distances, * p < 0.05; *** p < 0.001; n.s.: not significant.

We hypothesized that coupling between brain regions representing two neighbouring timescales increases when one timescale becomes unpredictive. Numerically, this can be expressed by setting the weights of neighbouring timescales to 1 and all other predictor weights to −1 (Figure 6B). This hypothesized pattern of weights was not matched by the weights observed for the LSTM (left: *p* = 0.83, *d* = 0.27; right: *p* = 0.348, *d* = 0.1; Euclidean distance compared to null distributions drawn from target BOLD activity shifted in time), which was expected given that the continuously updated timescales were not organized along a gradient in the first place. Critically, for sparsely updated timescales of the HM-LSTM, surprisal-modulated connectivity in the left hemisphere not only matched our hypothesis (left: *p* = 0.032, *d* = 0.5; right: *p* = 0.853, *d* = 0.23) but also matched our hypothesis better than the LSTM (left: *p* = 0.001, *d* = 0.79; right: *p* = 0.902, *d* = 0.3).

To specify the directionality of information flow, we averaged weights of top-down modulations (i.e., predictor weights of timescales longer than respective target timescales) and bottom-up modulations. We found no difference between the modulatory strength of these top-down and bottom-up connections (LSTM left: *p* = 0.184, *d* = 0.23; right: *p* = 0.839, *d* = 0.4; HM-LSTM left: *p* = 0.408, *d* = 0.4; right: *p* = 0.367, *d* = 0.16).

## Discussion

How are the complex temporal dependencies underlying natural speech processed in the brain to inform predictions of upcoming speech? In the present study, we emulated these prediction processes in two language models (artificial neural networks; LSTM vs. HM-LSTM), which critically differed in how often semantic context representations are updated at multiple, hierarchically organized timescales.

Surprisal as derived from both language models modulated reading times in the behavioural reading task to a similar degree, while hemodynamic brain responses to surprisal during the listening task differed between models: In early auditory cortex and supramarginal gyrus, the continuously updating LSTM predicted activity better than the sparsely updating HM-LSTM. In general, surprisal at the shortest timescale was decoded more precisely from temporo-parietal regions when derived from the continuously updating LSTM than from the HM-LSTM.

In contrast, and in line with our initial hypothesis, temporo-parietal regions hierarchically encoded the (sparsely updated) event-based surprisal provided by the layers of the HM-LSTM, with longer timescales represented in inferior parietal regions. Moreover, higher timescale-specific surprisal based on the HM-LSTM increased connectivity from receptive windows of a given timescale to their immediately neighbouring (shorter or longer) timescales.

Together, these results provide evidence for the neurobiological parsimony of an event-based processing hierarchy. In the present data, this was expressed in the simultaneous neural representation of surprisal at multiple timescales, and in surprisal dynamically gating the connectivity between these timescale-specific receptive windows.

### The event-based organization of context as a foundation for language prediction

The spatial organization of timescale-specific receptive windows observed in the present study converges with previous results, where bilateral primary auditory cortex coded for relatively shorter timescales (e.g., words) and inferior parietal cortex coded for longer timescales (e.g., paragraphs; Ref. ^28^). Critically, this spatial overlap was found despite targeting different aspects of speech processing.

In our study, neural responses were expressed as a function of timescale-specific word surprisal, a proxy tapping into prediction processes. In other studies, receptive windows were based on the (in-)consistency of neural activity across participants in response to speech input at varying timescales^50^, which is typically linked to working memory formation. This implies that the same neural system most likely fulfils distinct functions: Temporal receptive windows have been suggested to store timescale-specific context in working memory, and, in parallel, exploit this context to process information in the present^9^. In line with this theoretical account, our results suggest that timescale-specific memory representations serve as the basis for the generative models shaping predictions of upcoming speech.

The here observed temporo-parietal gradient of surprisal at sparsely updated representations of context is specifically well in line with accounts of neural event segmentation^51,52^, and with the notion of hierarchical multiscale network architectures more generally (here, HM-LSTM). Taking a sentence from our listening task as an example, “The wild wine was called ink grapes” is embedded in a brief event where the narrator describes how the bluish black of the grapes in a backyard reminded her of the color of the night. At the same time, the sentence is embedded in a larger event of the author wandering around the single rooms and the garden of her parents’ house, and an even larger event of the author reliving the memory of walking the streets in her Romanian hometown.

The HM-LSTM architecture resembles neural event segmentation in two decisive points. First, the boundary detector allows revealing the event structure of context, similar to an increase of neural activity indexing prediction errors at event boundaries^53-55^. Second, the sparse updates to higher processing stages at event boundaries allow retaining multiple, stable context representations in memory, similar to temporo-parieto-occipital receptive windows reflecting the hierarchical event structure during movie watching^36^. We directly tie in with this result by showing that such hierarchical, event-based context enables neural prediction processes.

What are the computational and mechanistic implications of this contextual architecture for the prediction of speech? Somewhat paradoxically, event models have been referred to as “an added burden for an organism”^56^. This argument is certainly plausible with regard to the size of the parameter space, which increases in an artificial or, likewise, biological neural network by introducing an additional boundary detector. At the expense of model parsimony, however, such an event-based network allows for less updates in comparison to continuously updating networks like the LSTM, where each new input to the model elicits computationally complex updates to all timescales.

The trade-off between computational costs of boundary detector versus update frequency is nicely illustrated by the fact that the sparsely updating HM-LSTM is considerably faster in making predictions than the continuously updating LSTM^42^. Thus, from a functional perspective, keeping layered representations of multiple events in memory allows to efficiently draw on diverse information to make predictions on upcoming speech.

### Context-dependent surprisal as a gating mechanism for predictions and prediction errors

The hallmark of prediction processes in our data is the increase in reading times and neural activity observed in response to more surprising input^19,57,58^. There are different computational ways in which this “expectation suppression”^59^ can be realized, namely integration difficulty, neural sharpening, and predictive coding.

One take on expectation suppression is that surprising sensory input is more difficult to integrate into already existing representations of context because it conveys a relatively larger amount of new information^60,61^. As the architecture of the sparsely updating HM-LSTM dictates that new information is integrated into timescale-specific representations only at event boundaries, integration difficulty should arise primarily at an event-by-event basis. However, surprisal indeed varies on a word-by-word basis. This conceptual mismatch renders it highly unlikely that integration difficulty accounts for our effects of event-based surprisal. The other two accounts both assume that expectation suppression is indicative of prediction processes but differ in how these processes are thought to be implemented in the brain.

Sharpening accounts argue that unexpected components of sensory input are suppressed via feedback predictions^62,63^, resulting in an overall decrease of neural activity in response to expected input. Under the predictive coding account^12,64^, the brain filters out (or “dampens”) expected components of sensory input, so remaining neural activity (mostly related to prediction errors) is overall smaller for more expected input. The similarity between hypothesized response patterns makes it notoriously hard to disentangle those accounts^32,63,65^.

Notably, however, a distinguishing feature of predictive coding is the specificity of feedforward prediction error signals, which can be captured by modelling effective connectivity between receptive windows of timescales. In agreement with the hierarchical information flow laid out in predictive coding^12^, surprisal in our study modulated connectivity via bidirectional links between neighbouring receptive windows of longer and shorter event-based timescales in the left hemisphere (Figure 6).

Surprisal in the event-based artificial neural network was modelled as the amount of information an input word conveys that cannot be explained away by the context (or generative model) represented at a specific timescale. Therefore, the increase of feedforward connectivity in response to higher surprisal precisely aligns with the concept of prediction errors in predictive coding^64^.

In addition, the increase of feedback connectivity in response to higher surprisal accords with an electrocorticography study in macaques by Chao and colleagues^22^. The study showed that prediction errors evoked in tone sequences trigger feedback signals from prefrontal to anterior temporal and early auditory cortex in alpha and beta frequency bands. Extending these previous results, our findings suggest that surprisal initiates bottom-up prediction errors, indicative of imprecise predictions, and top-down updates to predictions at processing stages of shorter events to facilitate perception of new words.

As an interim conclusion, our findings have two important implications for frameworks of prediction and prediction error: First, we show that a multi-layered hierarchy of predictive coding (e.g., Ref.^14^) applies well to higher-order semantic language processing. Second, predictive coding remains a viable account of neural processing, also when put to test using complex temporal dependencies underlying real-life stimuli.

### Implications for a larger network perspective on the event-based prediction hierarchy

Dual stream models of language propose that speech processing is organized along a ventral and a dorsal stream^66,67^. In the present study, we found a hierarchy of speech prediction along the dorsal stream, which emanated from early auditory cortex and extended well into parietal cortex (Figure 4B).

This result may seem at odds with other studies showing an additional mirror-symmetric ventral gradient, in which more complex speech features are represented in more anterior temporal regions^68^. The ventral stream has been proposed to chunk speech features into increasingly abstract concepts irrespective of their temporal presentation order^69^. In contrast, we here modelled context representations by respecting the temporal order of words, that is, the HM-LSTM integrates incoming words into an event until words become too dissimilar to previous words and a new event is created. Hence, the ventral stream may contribute to hierarchical speech prediction by exploiting another, more nested facet of context.

The inferior frontal gyrus (IFG), alongside premotor cortex, is deemed the apex of the dorsal stream^67^, yet we here considered only the role of temporo-parietal cortex in speech prediction. Previous studies showed that activity in IFG relies on longer timescales of speech being intact^28,70^, that connectivity between IFG and superior temporal gyrus is driven by expectations^71,72^, and that right IFG is sensitive to the violation of non-local regularities^73,74^. While this suggests an interplay between frontal and temporo-parietal regions in hierarchical speech prediction, the precise anticipatory mechanisms IFG exerts cognitive control over are just as unclear as how top-down cognitive control and bottom-up sensory input are balanced along the hierarchy.

Beyond short-term semantic context, also long-term knowledge facilitates speech prediction. In theory, both memory systems can be couched into the larger framework of the dual reference frame system^75^, where flexible sensory knowledge in parietal cortex interacts with stable conceptual knowledge in hippocampus. Consistent with the key characteristics of the speech prediction hierarchy, hippocampus codes for boundaries in the environment^76,77^, hierarchically organizes memories^78^ and engages in predictive coding^79,80^. As parietal cortex has been shown to interface with hippocampus at event boundaries of longer timescales during movie watching^36^, we speculate that the hierarchy of speech prediction might extend from receptive windows in parietal cortex to hippocampus.

Importantly, the event-based prediction hierarchy relies on a set of neural computations—i.e., event segmentation, temporal receptive windows, predictive coding—available beyond the domain of language. Our results thereby encourage future studies to probe its generalizability to other species, sensory modalities, and cognitive functions.

### Alternative mechanisms of predictive processing in lieu of event-based timescales

Although we only found a processing hierarchy for surprisal of sparsely updated timescales, temporo-parietal regions were nevertheless sensitive to continuously updated timescales. In particular, decoding accuracies suggested a predominance of the LSTM over the HM-LSTM at the shortest timescale and encoding accuracies suggested a predominance in medial temporal and anterior parietal regions (Figure 4C and 5B). This finding agrees with previous studies showing that participants track changes to situational dimensions of narratives both “globally” at the end of an event and “incrementally” within events^81^ and that computational models with continuous updates to all hierarchical levels can explain the construction of context representations in temporo-parietal regions^34^. Could continuously updated context representations, after all, play an integral role in successful speech prediction?

One potential explanation for the negligible neural effects at longer timescales is that the continuously updating language model relies primarily on shorter timescales in predicting the next word. This is supported by the considerably worse model performance observed for longer LSTM timescales (i.e., higher average word surprisal) compared to both shorter LSTM timescales and, despite comparable overall model performance, all HM-LSTM timescales (Supplementary Figure 1). Interestingly, the accuracy in predicting reading speed from word surprisal was the same between LSTM and HM-LSTM (Figure 2), suggesting that continuous updates make for an efficient mechanism to generate equally accurate predictions while relying on less timescales. The strength of such continuously updating models is that context representations are more integrated with what is currently relevant for prediction.

A unifying account might be that continuous and sparse updating mechanisms form one instead of two distinct processing streams. For example, Sainburg and colleagues^82^ showed that short-range dependencies of acoustic speech features follow sequential Markovian processes, whereas long-range dependencies follow hierarchical processes. This poses the question whether such interactions of different sequencing mechanisms also better match the semantic structure of speech. Future studies could test this by setting up hybrid language models with continuous updates on shorter and sparse updates on longer timescales.

## Conclusion

The present study bridges the gap between the hierarchical, temporally structured organization of context in language comprehension on the one hand and the more general principles of hierarchical predictive processing in cerebral cortex on the other hand.

Combining continuously narrated speech, artificial neural networks, and functional MRI building on these networks’ output allowed us, first, to sample the natural dynamic range of word-to-word changes in predictiveness over a multi-level hierarchy. Second, we were able to systematically compare the neural effects of different contextual updating mechanisms.

Our data demonstrate that the prediction processes in language comprehension build on an event-based organization of semantic context along the temporo-parietal pathway. Not least, we posit that such an event-based organization provides a blueprint for a semantically rich, yet computationally efficient network architecture of anticipatory processing in complex naturalistic environments.

## Materials and Methods

### Participants

Thirty-seven healthy, young students took part in the fMRI listening study. The final sample included *N* = 34 participants (18-32 years; *M* = 24.65; 18 female), as data from one participant was excluded from all analyses due to strong head movement throughout the recording (mean framewise displacement > 2 SD above group average^83^) and two experimental sessions were aborted because participants reported to not understand speech against noise. Another 26 students (19-32 years; *M* = 23.54; 17 female) took part in the behavioural self-paced reading study.

All participants were right-handed German native speakers who reported no neurological, psychiatric or hearing disorders. Participants gave written informed consent and received an expense allowance of €10 per hour of testing. The study was conducted in accordance with the Declaration of Helsinki and was approved by the local ethics committee of the University of Lübeck.

### Stimulus materials

As a speech stimulus in the fMRI listening task, we used the first 64 minutes of an audio recording featuring Herta Müller, a Nobel laureate in Literature, reminiscing about her childhood as part of the German-speaking minority in the Romanian Banat (“Die Nacht ist aus Tinte gemacht”, 2009).

To emulate an acoustically challenging scenario in which listeners are likely to make use of the semantic predictability of speech^84^, this recording was energetically masked by a stream of concatenated five-second sound textures at a signal-to-noise ratio of 0 dB. Sound textures were synthesized from the spectro-temporal modulation content of 192 natural sounds (i.e., human and animal vocalizations, music, tools, nature scenes^85^), so that the noise stream did not provide any semantic content potentially interfering with the prediction of upcoming speech. The order in which sound textures were arranged was randomized across participants. For more details on how sound textures in the present experiment were generated and how they were processed in auditory cortex, see Ref.^40^.

The monaural speech and noise streams were sampled to 44.1 kHz and custom filters specific to the left and right channel of the earphones used in the fMRI experiment were applied for frequency response equalization. Finally, speech-in-noise stimuli were divided into eight excerpts à 8 minutes, which served as independent runs in the experiment.

A trained human external agent literally transcribed the speech stream. The text transcript comprised 9,446 words, which were used as stimuli in the self-paced reading task and as input to our language models. To automatically determine onset and offset times of all spoken words and phonemes, we used the web service of the Bavarian Archive for Speech Signals (BAS)^86^: First, the text transcript was transformed to a canonical phonetic transcript encoded in SAM-PA by the G2P module. Second, the most likely pronunciation for the phonetic transcript was determined within a Markov model and aligned to the speech recording by the MAUS module. Fourteen part-of-speech tags were assigned to the words in the text transcript using the pre-trained German language model de_core_news_sm (2.2.5) from spaCy (https://spacy.io/). Based on these tags, words were classified as content or function words. Word frequencies were derived from the subtitle-based SUBTLEX-DE corpus^87^ and transformed to standardized Zipf values^88^ operating on a logarithmic scale from about 1 (word with a frequency of 1 per 100 million words) to 7 (1 per 1,000 words). The Zipf value of a word not observed in the corpus was 1.59 (i.e., smallest possible value).

### Experimental procedures

#### Behavioural self-paced reading task

While the transcribed story was presented word-by-word on a noncumulative display, participants had the task to read each word once at a comfortable pace and quickly press a button to reveal the next word as soon as they had finished reading. A timeout of 6 seconds was implemented. The time interval between word appearance and button press was logged as the reading time. After each run, participants answered three four-option multiple-choice questions on the plot of the story (performance: *Ra* = 58.33-100 % correct, *M* = 79.17, *SD* = 10.87) and took a self-paced break. In total, each participant completed four out of eight runs, which were randomly selected and presented in chronological order. Throughout the reading task, we recorded movement and pupil dilation of participants’ right eye at a sampling rate of 250 Hz in one continuous shot with an eye tracker (EyeLink 1000, SR Research).

The experiment was controlled via the Psychophysics Toolbox^89^ in MATLAB (R2017b, MathWorks). All words were presented 20 % off from the left edge of the screen in white Arial font on a grey background with a visual angle of approximately 18°. Participants used a response pad (URP48, The Black Box Toolkit) to navigate the experiment with their right index finger. The experimental session took approximately 40 minutes.

#### Functional MRI listening task

We instructed participants to carefully listen to the story while ignoring the competing stream of sound textures as well as the MRI scanner noise in the background. Each of the eight runs was initialized by 10 baseline MRI volumes after which a white fixation cross appeared in the middle of a grey screen and playback of the 8-minute audio recording started. MRI recording stopped with the end of playback and participants successively answered the same questions used in the self-paced reading task via a response pad with four buttons (HHSC-2×4-C, Current Designs). On average, participants answered 65.5 % of the questions correctly (*Ra* = 38-100 %, *SD* = 15.9 %). There was a 20-second break between consecutive runs.

The experiment was run in MATLAB (R2016b) using the Psychophysics Toolbox. Stimuli were presented at a subjectively comfortable sound pressure level via insert earphones (S14, SENSIMETRICS) covered with circumaural air cushions. The experimenters monitored whether participants kept their eyes open throughout the experiment via an eye tracker.

#### MRI data acquisition

MRI data were collected on a 3 Tesla Siemens MAGNETOM Skyra scanner using a 64-channel head coil. During the listening task, continuous whole-brain fMRI data were acquired in eight separate runs using an echo-planar imaging (EPI) sequence (repetition time (TR) = 947 ms, echo time (TE) = 28 ms, flip angle = 60°, voxel size = 2.5 × 2.5 × 2.5 mm, slice thickness = 2.5 mm, matrix size = 80 × 80, field of view = 200 × 200 mm, simultaneous multi-slice factor = 4). Fifty-two axial slices were scanned in interleaved order. For each run, 519 volumes were recorded.

Before each second run, field maps were acquired with a gradient echo (GRE) sequence (TR = 610 ms, TE_1_ = 4.92 ms, TE_2_ = 7.38 ms, flip angle = 60°, voxel size = 2.5 × 2.5 × 2.75 mm, matrix size = 80 × 80, axial slice number = 62, slice thickness = 2.5 mm, slice gap = 10%).

In the end of an experimental session, anatomical images were acquired using a T1-weighted (T1w) MP-RAGE sequence (TR = 2,400 ms, TE = 3.16 ms, flip angle = 8°, voxel size = 1 × 1 × 1 mm, matrix size = 256 × 256, sagittal slice number = 176) and a T2-weighted (T2w) SPACE sequence (TR = 3,200 ms, TE = 449 ms, flip angle = 120°, voxel size = 1 × 1 × 1 mm, matrix size = 256 × 256, sagittal slice number = 176).

### Modelling the predictiveness of context at multiple timescales

We trained two versions of a long short-term memory network (LSTM) with five layers to predict the next word in a story given a sequence of semantic context: a “continuously updating LSTM” where information is fed to a higher layer with each upcoming word, and a competing “sparsely updating HM-LSTM” where information is fed to a higher layer only at the end of a timescale. The predictiveness of context at multiple timescales was read out from single layers of both language models for each word in the story presented to participants in experiments. Ultimately, we tested how closely these derivatives of different network architectures match signatures of behavioural and neural prediction processes.

#### Representing words in vector space

In natural language processing, it is common to represent a word by its linguistic features in the form of high-dimensional vectors (or embeddings). As the German language is morphologically rich and flexibly combines words into new compounds, there are many rare words for which language models cannot learn good (if any) vector representations on the word level. Therefore, we mapped all texts used for training, validating and testing our language models to pre-trained *sub*word vectors publicly available in the BPEmb collection^90^. These embeddings allow for the representation of *any* word by a combination of 100-dimensional subwords from a finite vocabulary of 100,000 subwords. We further reduced this vocabulary to those subwords that appeared at least once in any of the texts used for training, validating or testing our language models (i.e., number of subwords in vocabulary v= 91,645). See Supplementary Text 1 for a detailed description of the BPEmb vocabulary.

Matching our texts to subwords and their respective embeddings in the BPEmb vocabulary, yielded the embedded text t ∈ *R*^*w*×*e*^ where *w* is the number of word sand *e* = 100 is the number of vector dimensions. On average, a word in the story was represented by 1.07 subwords (*Ra* = 1-6, *SD* = 0.33). As single words were encoded by only one subword in 94.25 % of cases, we will refer to subwords as words from here on.

#### Architecture of language models

When listening to a story, a fused representation of all spoken words {*w*_1_, *w*_2_,…, *w_p_*} is maintained in memory and used as context information to make a prediction about the upcoming word *w*_*p*+1_. In natural language processing, this memory formation is implemented via recurrent connections between the states of adjacent neural network cells. The hidden state *h*_*p*-1_ stores all relevant context and is sequentially passed to the next cell where it is updated with information from word *w_p_*.

As such a simple recurrent neural network (RNN) tends to memorize only the most recent past, the more complex LSTM^41^ became a standard model in time series forecasting. In an LSTM cell, the state is split in two vectors: The cell state *c_p_* acts as long-term memory, whereas the hidden state *h_p_* incorporates information relevant to the cell output (i.e., the prediction of the next word). The integration of new information and the information flow between the two memory systems is controlled by three gating mechanisms.

When stacking multiple LSTM cells on top of each other, semantic context gets hierarchically organized in the model, with lower layers coding for short-term dependencies and higher layers coding for long-term dependencies between words. The bottom-up input to the first layer remains to be the embedded word *w_p_*. However, the lower layer’s hidden state 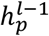 becomes the input to a cell from the second layer on. Importantly, the hidden state and cell state are updated at each layer with every new bottom-up input to the model.

A competing model that has been shown to slightly outperform the continuously updating (“vanilla”) LSTM in character-level language modelling is the hierarchical multiscale LSTM (HM-LSTM)^42^. This model, referred to as “sparsely-updating HM-LSTM”, employs a revised updating rule where information from the lower layer is only fed forward at the end of a timescale (i.e., a sequence of words closely related to each other).

Importantly, The HM-LSTM allows for a sparse updating rate, with lower layers operating on short timescales and higher layers operating on longer timescales. Here, we used the simplified version of the HM-LSTM^91^ with no top-down connections. See Supplementary Text 2 for a detailed description of the model architecture including all relevant formulas.

#### Prediction of the next word

LSTM and HM-LSTM cells form the representations of semantic information relevant to speech prediction, whereas the actual prediction of the next word takes place in the output module. Here, hidden states at word position *p* are combined across the different layers of the language model. The combined hidden state 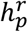 is mapped to a fully connected dense layer of as many neurons as there are words in the vocabulary and squashed to values in the interval [0,1], which sum to 1 (i.e., softmax function). Each neuron in resulting vector *d_p_* indexes one particular word in vocabulary *v* and denotes its probability of being the next word. Finally, the word referring to the highest probability in the distribution is chosen as the predicted next word *s_p_* in a story. See Supplementary Text 3 for a detailed description of word prediction including all relevant formulas.

#### Training and evaluation of language models

The objective of our language models was to minimize the difference between the “predicted” probability distribution *d_p_* (i.e., a vector of probabilities ranging from 0 to 1) and the “actual” probability distribution corresponding to the next word in a text (i.e., a vector of zeros with a one-hot encoded target word). To this end, we trained models on mini-batches of 16 independent text sequences à 500 words and monitored model performance by means of categorical cross-entropy between the “predicted” and “actual” probability distribution of each word in a sequence. Based on model performance, trainable parameters were updated after each mini batch using the Adam algorithm for stochastic gradient optimization^92^.

Our text corpus comprised more than 130 million words including 4,400 political speeches^93^ as well as 1,200 fictional and popular scientific books. All texts had at least 500 words; metadata, page numbers, references and punctuations (except for hyphenated compound words) were removed from documents. A held-out set of 10 randomly selected documents was used for validation after each epoch of training (i.e., going through the complete training set once) and allowed us to detect overfitting on the training set. Training automatically stopped after model performance did not increase over two epochs for the validation data set.

Using a context window of 500 words, we aimed at roughly modelling timescales of the length of common linguistic units in written language (i.e., words, phrases, sentences, and paragraphs). Therefore, we only used a small range of values from three to seven to find the number of layers— intended to represent distinct timescales—best suited to make good predictions. Additionally, we tuned the number of units in LSTM and HM-LSTM cells of language models, using values from 50 to 500 in steps of 50. Hyperparameters were evaluated on a single epoch using grid search and the best combination of hyperparameters was chosen based on performance on the validation set. Our final language models had five LSTM or HM-LSTM layers à 300 units and an output module. The LSTM model had 31,428,745 and the HM-LSTM model had 31,431,570 trainable parameters. Models were trained and evaluated with custom scripts in TensorFlow 2.1^94^. See Supplementary Text 4 for a detailed description of architectural choices.

#### Deriving the predictiveness of timescales by “lesioning” the language models

We used each trained language model to determine the predictiveness of semantic context in the story presented to participants in the behavioural and fMRI experiment. First, predictiveness was read out from “full” models: We iteratively selected each word in the story as a target word and fed all 500 context words preceding the target word to our language models. Note that the context for target words in the very beginning of the story comprised less than 500 words. The “predicted” probability of each word in the vocabulary was extracted from distribution d_p_ in the output module.

Second, predictiveness was read out from “lesioned” models, where we allowed information to freely flow through networks, yet only considered semantic context represented at single layers to generate the “predicted” probability distribution. These timescale-resolved probabilities were created by setting weight matrix 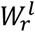 of pre-trained models to zero for all layers of no interest, so that the hidden state of only one layer is passed to the softmax function and all other layers have no bearing on the final prediction. We iteratively set all but one layer to zero with each layer being the only one influencing predictions once, resulting in five lesioned outputs for each language model.

We derived three measures of predictiveness from probability distributions. Our primary measure was the degree of surprisal associated with the occurrence of a word given its context. Word surprisal is the negative logarithm of the probability assigned to the actual next word in a story.

Secondary measures of predictiveness were used to explore the specificity of the processing hierarchy to only some aspects of prediction processes. Word entropy reflects the amount of uncertainty across the whole probability distribution, which is the negative sum of probabilities multiplied by their natural logarithm. When high probabilities are assigned to only one or few words in the vocabulary, entropy is low. On the other hand, entropy is high when semantic context is not informative enough to narrow predictions down to a limited set of words, resulting in similar probabilities for all candidate words. As all information necessary to determine the entropy of a word is already available to participants before word presentation, entropy of word *w_p_* was ascribed to the previous word *w*_*p*-1_. Whereas word surprisal quantifies the availability of information on the actual next word, word entropy quantifies the overall difficulty of making any definite prediction. Another secondary measure of predictiveness was the relatedness of the predicted next word to the actual next word. This word similarity is expressed as the correlation of respective word embeddings.

A high positive Product-moment correlation indicates that the prediction is semantically close to the target word, even though the model prediction might have been incorrect.

All three measures were calculated for each word in the story, separately for full models and five lesioned models. This yielded an 18-dimensional feature space of predictiveness for the LSTM as well as the HM-LSTM model, which was linked to BOLD activity and reading times in our analysis.

Additionally, we created a metric to dissociate neural effects of predictiveness from more low-level effects of semantic dissimilarity between target words and their preceding context. To this end, we correlated the embedding of each function word in the story with the average embedding of a context window, and subtracted resulting Product-moment correlation coefficients from 1^95^. This measure of contextual dissimilarity was calculated at five timescales corresponding to a logarithmic increase in context length (i.e., 2, 4, 8, 16, and 32 words).

To determine temporal integration windows of layers, we scrambled input to language models at nine levels of granularity corresponding to a binary logarithmic increase in the length of intact context (i.e., context windows of 1-256 words). For each layer, we fit linear functions to word surprisal across context windows and extracted slope parameters indicating how much a layer benefits from longer context being available when predicting the next word. On the second level, we fit linear functions to these layer-specific integration windows to determine the context benefit of higher layers over shorter layers. Resulting model-specific slopes were compared to a null distribution of slopes computed by shuffling integration windows across layers (n = 10,000). Additionally, slopes were compared between language models by means of a Monte Carlo approximated permutation test (n = 10,000) on the difference of means.

#### Convolving features with the hemodynamic response function

We used three classes of features to model brain responses: 18 features of the predictiveness of timescales (per language model), 3 linguistic features, and 9 acoustic features. While we were primarily interested in modelling effects of predictiveness, linguistic and acoustic features were used as nuisance regressors potentially covarying with predictiveness. Linguistic features included information on when words were presented (coded as 1), whether they were content or function words (coded as 1 and −1), and which frequency they had. In Ref. ^40^, we decomposed the speech-in-noise stimuli into a 288-dimensional acoustic space of spectral, temporal and spectro-temporal modulations, which was derived from a filter bank modelling auditory processing^96^. Here, we reduced the number of acoustic features to the first 9 principal components, which explained more than 80 % of variance in the original acoustic space. All features were z-scored per run.

A set of 500 scrambled features of predictiveness was generated, which was used to estimate null distributions of predictive processing. We applied the fast Fourier transform to single features, randomly shifted the phase of frequency components, and inverted the transform to project the data back into the time domain. This preserved power spectra of features but disrupted the temporal alignment of frequencies. See Supplementary Text 5 for a detailed description of convolving features with the hemodynamic response function (HRF).

### Data analysis

See Supplementary Text 6 for a detailed description of structural and functional MRI data preprocessing.

#### Selection of regions of interest

We hypothesized that the speech prediction hierarchy is represented as a gradient along a temporo-parietal pathway. This rather coarse region of interest was further refined to only include regions implicated in speech processing. To this end, we used intersubject correlation^45^ as a measure of neural activity consistently evoked across participants listening to speech in noise. As we were primarily interested in shared responses to the speech stream, this approach allowed us to leverage the inconsistency of the noise stream across participants. The presentation of sound textures in different order likely evoked more heterogeneous neural responses, leading to a diminished shared representation of the noise stream. Therefore, we inferred that the shared neural responses we observed were largely driven by the speech stream, which was the same for all participants.

At the first level, hyperaligned functional time series of each participant (see Supplementary Text 6) were concatenated across experimental runs and correlated with every other participant on a vertex-by-vertex basis, resulting in pairwise maps of intersubject Product-moment correlations. A group map was created by calculating the median correlation coefficient across pairs of participants for each vertex. At the second level, we applied a bootstrap hypothesis test with 10,000 iterations. To create the null distribution, we iteratively resampled participants with replacement and derived median group maps from their pairwise correlation maps. When the same participant was sampled more than once in a bootstrap iteration, the pairwise correlation map of that participant with herself was not included in the computation of the group map. The actual median intersubject correlation was ranked against the normalized null distribution to obtain a p-value for each vertex. Intersubject correlations were computed with the Python package BrainlAK^97^.

Finally, we used a multi-modal parcellation^46^ to select those lateral temporal and parietal parcels of which at least 80% of the vertices had a significant intersubject correlation (*p* < 0.01, adjusted for false discovery rate; FDR)^98^ in one hemisphere. The following parcels were included in the region of interest (ROI): early auditory cortex (EAC), auditory association cortex (AAC), lateral temporal cortex (LTC), temporo-parietal-occipital junction (TPOJ), inferior parietal cortex (IPC), and superior parietal cortex (SPC). As the temporal MT+ complex is thought to be mainly involved in visual processing, this region was not considered an appropriate candidate parcel. All further analyses including MRI data were limited to the temporo-parietal ROI, which was organized along the anterior-posterior (left: 124 mm, right: 167 mm) and inferior-superior axis (left: 234 mm, right: 212 mm).

#### Functional data analysis

The starting point of our analyses was the question whether the timescales of speech prediction organize along a temporo-parietal processing hierarchy. In a forward model, we encoded the predictiveness of timescales into univariate neural activity and fit a gradient along the peak locations sensitive to specific timescales of surprisal. Next, we compared the explanatory power of both language models in a backward model, which decoded surprisal at different timescales from multivariate patterns of neural activity in temporo-parietal parcels. Finally, we modelled functional connectivity between peak locations to test whether the timescales of surprisal gate the information flow along the gradient.

##### Encoding model

The encoding approach (similar to e.g., Ref.^99^) allowed us to quantify for each temporo-parietal vertex, which features of predictiveness it preferentially represents. Two separate encoding models were estimated for each vertex in the ROI of single participants, one for each language model. Besides the features of predictiveness specific to language models, both models included the same linguistic and acoustic features as nuisance regressors. We modelled neural activity as a function of 30 HRF-convolved features characterizing speech and noise stimuli by:

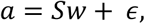

where *a*^*samples*×1^ is the activity vector (or BOLD time course) corresponding to a vertex, *S^samples×features^* is the stimulus matrix of features, *w*^*features*×1^ is a vector of estimated model weights, and *ϵ*^*samples*×1^ is a vector of random noise.

All models were estimated using ridge regression with four-fold cross validation. We paired odd-numbered functional runs with their subsequent even-numbered run, resulting in four data splits per participant. Each of the four data splits was selected as a testing set once; all other data splits were used as a training set. Within each fold, generalized cross-validation^100^ was carried out on the training set to find an optimal estimate of regularization parameter λ from the data, searching 100 values evenly spaced on a logarithmic scale from 10^-5^ to 10^8^. Weights of predictiveness were extracted from the model fit with the optimal regularization parameter and averaged across cross-validation folds to obtain stable weights.

To evaluate the performance of encoding models and their ability to generalize to new data, we applied the weights estimated on the training set to the features of the held-out testing set in each cross-validation fold. The predicted BOLD time series was correlated with the actual BOLD time series. The resulting Product-moment correlation coefficient is the encoding accuracy, which was averaged across cross-validation folds and Fisher z-transformed.

Additionally, we created null distributions of weights and encoding accuracies by estimating forward models on scrambled features of predictiveness (similar to e.g., Ref. ^101^). We set up 500 separate models, which included scrambled features of predictiveness but intact linguistic and acoustic features. Models were estimated largely following the cross-validation scheme outlined for observed data. However, we re-used optimal regularization parameters from non-scrambled models of corresponding folds. All ridge regression models were implemented using the RidgeCV function in the Python package scikit-learn^102^.

##### Peak selection

For both language models, we derived five temporo-parietal maps in the left and right hemisphere of single participants: one weight map for each timescale of word surprisal. Maps represented the sensitivity of brain regions to timescale surprisal; positive weights indicate increasing BOLD activity to more surprising words.

To illustrate the location and extent of brain regions modulated by timescale surprisal, we performed an analysis similar to cluster-based permutation tests in Fieldtrip^103^. For each timescale, vertex-wise weights observed across participants were tested against zero by means of a one-sample *t*-test. We combined a vertex into a cluster with its adjacent vertices if it was significant at an alpha level of 0.05 and had at least two significant neighbours. We clustered vertices with negative *t*-values separately from vertices with positive *t*-values. The summed *t*-value of an observed cluster served as the cluster-level statistic and was compared with a Monte Carlo approximated null distribution of summed *t*-values. This null distribution was created by performing clustering on scrambled partitions of timescale-specific weight maps and selecting the largest summed *t*-value for each partition. An observed cluster was considered significant if its summed *t*-value was exceeded by no more than 2.5 % of summed *t*-values from scrambled partitions.

Beyond this rather coarse mapping of temporo-parietal brain regions onto the timescales of surprisal, our main analysis focused on how timescale-specific peak locations distribute along the inferior-superior axis only. Of note, we hypothesized that a hierarchy of speech prediction evolves from temporal to parietal areas, which corresponds to the inferior-superior axis of our ROI. A window with a height of 2 mm was shifted along the inferior-superior axis of the temporo-parietal ROI in steps of 1 mm. All weights of a timescale falling into the window were averaged, thereby collapsing across the anterior-posterior axis. The resulting one-dimensional weight profile of a timescale spanned inferior to superior locations and was smoothed using robust linear regression over a window of 70 mm. For each unilateral weight profile of single participants, local maxima (i.e., a sample larger than its two neighbouring samples) were determined.

We applied two different approaches to select one peak location for each timescale from these local maxima. In the naïve approach of peak selection, the local maximum with the highest positive value was defined as a peak. As this approach makes it hard to find a consistent order of timescales when surprisal is not just processed along the dorsal but also the ventral processing stream, we also applied a pre-informed peak selection approach explicitly targeting the dorsal stream. Here, the peak of the first timescale had to be in the inferior half of the axis (i.e., temporal regions) and peaks of longer timescales had to be superior to the peak of the first timescale. Whenever no timescale peak could be defined, the largest positive value was selected. Both peak selection approaches yielded five timescale-specific coordinates on the inferior-superior axis for each participant, hemisphere and language model.

Additionally, we applied naïve peak selection to weight maps, which were rotated by −45° before collapsing across the first dimension. In the left hemisphere, an increase on that new axis indicated a shift to more superior and posterior regions, thereby simultaneously modelling effects on the inferior-superior and anterior-posterior axis. However, original right-hemispheric maps already had a strong rotation off the inferior-superior axis, thus rotating these maps rather brought them into alignment with the inferior-superior axis of non-rotated left-hemispheric maps.

##### Gradient fitting

We fit linear functions to coordinates of single participants across the timescales of surprisal. Models included an intercept term and the slope parameter was extracted from each fit. A positive slope indicates a gradient of timescale surprisal, where shorter timescales are represented in more inferior (anterior) temporo-parietal regions than longer timescale, which are represented in more superior (posterior) regions. We tested grand-average slope parameters against a null distribution of slopes with 10,000 partitions, which was created by randomly shuffling the coordinates of single participants across the timescales of surprisal and recalculating their slopes. As the first timescale was pre-set to have the most inferior coordinate in the pre-informed peak selection approach, this specific coordinate was not shuffled when calculating the null distribution for this approach. To compare slope parameters between language models, we performed a Monte Carlo approximated permutation test (n = 10,000), using the difference of means as a test statistic. As secondary analyses, gradients of predictive processing were also calculated for the timescales of word entropy and similarity. In a control analysis, a gradient was fit to timescale peaks following the same procedure described above but replacing features of predictiveness by contextual dissimilarity when estimating forward models.

To round off the encoding analysis, we compared temporo-parietal encoding accuracies between both language models. As we were interested in effects specific to the predictiveness of speech, encoding accuracies were z-scored to the null distribution of accuracies from scrambled features of predictiveness. A cluster-based permutation paired-sample *t*-test was calculated (n = 1,000, vertex-specific alpha level: 0.05, cluster-specific alpha level: 0.05). In comparison to the cluster test described above for the weight maps, we here created a null distribution of summed *t*-values by contrasting accuracies of language models whose labels had been randomly shuffled in single participants.

##### Decoding model

In our decoding approach (similar to e.g., Ref. ^99^), we quantified how much information multiple vertices jointly contain about a feature of predictiveness. For each language model, five separate backward models were estimated in each of six temporo-parietal parcels of single participants, one for each timescale of word surprisal. We modelled timescale-specific word surprisal as a function of neural activity in all vertices forming a parcel by:

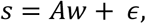

where *S*^*samples*×1^ is the stimulus vector of a feature, *A*^*samples×vertices*^ is the activity matrix of BOLD time courses corresponding to the vertices of a parcels, *w*^*vertices*×1^ of model weights, *ϵ*^*samples*×1^ is a vector of random noise.

The same cross-validation scheme as described for the encoding model was applied. However, instead of predicting BOLD activity, we here reconstructed surprisal at different timescales. By correlating the actual stimulus time series with the one predicted on the held-out testing set, we obtained the decoding accuracy of a parcel. Decoding accuracies were z-scored to the null distribution of accuracies determined for scrambled stimulus time series. We compared decoding accuracies between language models in each hemisphere, parcel and timescale by means of a Monte Carlo approximated permutation test (n = 10,000) on the difference of means. Resulting *p*-values were corrected for multiple comparisons using FDR correction.

##### Functional connectivity

To model the information flow between brain regions sensitive to the different timescales of word surprisal, we identified five unique seeds for both language models in each temporo-parietal hemisphere. On the inferior-superior axis, we re-used the grand-median coordinate of each timescale as localized in the pre-informed peak selection. The corresponding coordinate on the anterior-posterior axis was localized by shifting a moving average with a window centred on the inferior-superior coordinate along the anterior-posterior axis (width: 2 mm, height: 5 mm), and determining peak locations on smoothed weight profiles of single participants. Following, we placed a sphere with a radius of 5 mm on peak coordinates from both axes and averaged BOLD time courses of vertices falling within this sphere, yielding the timescale-specific neural activity of seeds.

We expected increased information flow between seeds of adjacent timescales when one timescale becomes uninformative for the prediction of upcoming speech. This modulatory influence of surprisal on connectivity was modelled along the lines of a psychophysiological interaction (PPI)^104^. In a standard PPI analysis, the neural time series of one brain region is regressed onto the pointwise product of an experimental stimulus and the neural time series of another brain region. Here, we extended this approach by creating timescale-specific interactions: BOLD time series of seeds were multiplied by their corresponding HRF-convolved surprisal time series but not any of the surprisal time series at another timescale.

Functional connectivity was calculated for both language models in each participant and hemisphere. We set up five regression models, with every seed being selected as a target once. The physiological (BOLD) time series of the target seed was mapped onto the physiological, psychological (timescale-specific surprisal), and psychophysiological time series from all other (predictor) seeds. Models were estimated within the same cross-validation scheme outlined for the encoding model. We extracted all four weights from psychophysiological interaction terms of each target seed and arranged weights in a 5-by-5 matrix, with target seeds on the main diagonal and predictor seeds off the diagonal. This matrix of observed psychophysiological interactions was compared to a matrix with hypothesized interaction weights: The diagonals below and above the main diagonal were set to 1 (indicating increased coupling when surprisal at a neighbouring timescale is high), all other items were set to −1. We calculated the Euclidean distance of single-participant matrices to this hypothesized matrix. The mean of observed Euclidean distances was compared to a null distribution of 10,000 mean Euclidean distances calculated on BOLD time series of target seeds randomly shifted in time by the number of samples in 1 to 7 functional runs. Euclidean distances were compared between language models in each hemisphere by means of a Monte Carlo approximated permutation test (n = 10,000) on the difference of means.

#### Behavioural data analysis

Reading times were used to test the behavioural relevance of the predictiveness determined by our language models. Trials with reading times shorter than 0.001 seconds or longer than 6 seconds were considered invalid and excluded. Further, we converted reading times to speed (number of words per 100 seconds) and excluded trials exceeding 3 standard deviations within a run and participant from all further analyses. On average, 1.31 % of trials (*SD* = 1.12, *Ra* = 0.32-6.15) were removed. Finally, reading speed was z-scored within runs.

For each participant, we predicted reading speed in a forward model, adopting the same cross-validated ridge regression scheme used for the analysis of fMRI data. Our feature space included the predictiveness of words as well as word frequency, word length (number of letters), content vs. function words and trial number as nuisance regressors. As this was a high-pace task, some features might have unfolded their effect on reading speed only over the course of a few words. Therefore, we added time-lagged versions of features to the model, that is, shifting features by −2 to 5 word positions. There were no lagged versions of the predictor coding for trial number added to the model.

To investigate whether predictiveness had an effect on reading speed beyond the effect of nuisance regressors, we compared the predictive accuracy of forward models in single participants to a null distribution of accuracies from models with scrambled features of predictiveness. The performance of language models was compared by z-scoring observed encoding accuracies to the null distribution and running a Monte Carlo approximated permutation test (n = 10,000) on the difference of means. This analysis was also carried out for the timescales of contextual dissimilarity.

## Acknowledgements

We thank Anne Herrmann, Clara Mergner, Malte Naujokat, Anna Ruhe, and Svenja Meyn for their help with data acquisition; Christine Sickert for her help in preparing the text corpus; Martin Göttlich for setting up the MR sequences; Mattias Heinrich for discussions on natural language processing; and Malte Wöstmann for suggestions to the reading-task design.

## Data availability

All functional data are publicly available on the Open Science Framework (OSF; https://osf.io/zbuah).

## References

1. Clark, A. Whatever next? Predictive brains, situated agents, and the future of cognitive science. Behavioral and Brain Sciences 36, 181–204 (2013).

2. Murray, J. D. et al. A hierarchy of intrinsic timescales across primate cortex. Nat Neurosci 17, 1661–1663 (2014).

3. Zhang, W. & Yartsev, M. M. Correlated Neural Activity across the Brains of Socially Interacting Bats. Cell 178, 413–428 (2019).

4. La Camera, G. et al. Multiple Time Scales of Temporal Response in Pyramidal and Fast Spiking Cortical Neurons. Journal of Neurophysiology 96, 3448–3464 (2006).

5. Burt, J. B. et al. Hierarchy of transcriptomic specialization across human cortex captured by structural neuroimaging topography. Nat Neurosci 21, 1251–1259 (2018).

6. Lakatos, P. et al. An Oscillatory Hierarchy Controlling Neuronal Excitability and Stimulus Processing in the Auditory Cortex. Journal of Neurophysiology 94, 1904–1911 (2005).

7. Mattar, M. G., Kahn, D. A., Thompson-Schill, S. L. & Aguirre, G. K. Varying Timescales of Stimulus Integration Unite Neural Adaptation and Prototype Formation. Current Biology 26, 1669–1676 (2016).

8. Lamme, V. A. F. & Roelfsema, P. R. The distinct modes of vision offered by feedforward and recurrent processing. Trends in Neurosciences 23, 571–579 (2000).

9. Hasson, U., Chen, J. & Honey, C. J. Hierarchical process memory: memory as an integral component of information processing. Trends in Cognitive Sciences 19, 304–313 (2015).

10. Buracas, G. T., Zador, A. M., DeWeese, M. R. & Albright, T. D. Efficient Discrimination of Temporal Patterns by Motion-Sensitive Neurons in Primate Visual Cortex. Neuron 20, 959–969 (1998).

11. Runyan, C. A., Piasini, E., Panzeri, S. & Harvey, C. D. Distinct timescales of population coding across cortex. Nature 548, 92–96 (2017).

12. Friston, K. A theory of cortical responses. Phil. Trans. R. Soc. B 360, 815–836 (2005).

13. Keller, G. B. & Mrsic-Flogel, T. D. Predictive Processing: A Canonical Cortical Computation. Neuron 100, 424–435 (2018).

14. Kiebel, S. J., Daunizeau, J. & Friston, K. J. A Hierarchy of Time-Scales and the Brain. PLoS Computational Biology 4, e1000209 (2008).

15. Chaudhuri, R., Knoblauch, K., Gariel, M.-A., Kennedy, H. & Wang, X.-J. A Large-Scale Circuit Mechanism for Hierarchical Dynamical Processing in the Primate Cortex. Neuron 88, 419–431 (2015).

16. Demirtaş, M. et al. Hierarchical Heterogeneity across Human Cortex Shapes Large-Scale Neural Dynamics. Neuron 101, 1181–1194.e13 (2019).

17. Bastos, A. M. et al. Visual Areas Exert Feedforward and Feedback Influences through Distinct Frequency Channels. Neuron 85, 390–401 (2015).

18. Cocchi, L. et al. A hierarchy of timescales explains distinct effects of local inhibition of primary visual cortex and frontal eye fields. eLife 5, e15252 (2016).

19. Wacongne, C. et al. Evidence for a hierarchy of predictions and prediction errors in human cortex. Proceedings of the National Academy of Sciences 108, 20754–20759 (2011).

20. Schwiedrzik, C. M. & Freiwald, W. A. High-Level Prediction Signals in a Low-Level Area of the Macaque Face-Processing Hierarchy. Neuron 96, 89–97.e4 (2017).

21. Donhauser, P. W. & Baillet, S. Two Distinct Neural Timescales for Predictive Speech Processing. Neuron 105, 385–393.e9 (2020).

22. Chao, Z. C., Takaura, K., Wang, L., Fujii, N. & Dehaene, S. Large-Scale Cortical Networks for Hierarchical Prediction and Prediction Error in the Primate Brain. Neuron 100, 1252–1266.e3 (2018).

23. Honey, C. J. et al. Slow Cortical Dynamics and the Accumulation of Information over Long Timescales. Neuron 76, 423–434 (2012).

24. Stephens, G. J., Honey, C. J. & Hasson, U. A place for time: the spatiotemporal structure of neural dynamics during natural audition. Journal of Neurophysiology 110, 2019–2026 (2013).

25. Mesgarani, N., Cheung, C., Johnson, K. & Chang, E. F. Phonetic Feature Encoding in Human Superior Temporal Gyrus. Science 343, 1006–1010 (2014).

26. Huth, A. G., de Heer, W. A., Griffiths, T. L., Theunissen, F. E. & Gallant, J. L. Natural speech reveals the semantic maps that tile human cerebral cortex. Nature 532, 453–458 (2016).

27. de Heer, W. A., Huth, A. G., Griffiths, T. L., Gallant, J. L. & Theunissen, F. E. The Hierarchical Cortical Organization of Human Speech Processing. The Journal of Neuroscience 37, 6539–6557 (2017).

28. Lerner, Y., Honey, C. J., Silbert, L. J. & Hasson, U. Topographic Mapping of a Hierarchy of Temporal Receptive Windows Using a Narrated Story. Journal of Neuroscience 31, 2906–2915 (2011).

29. Bornkessel-Schlesewsky, I., Schlesewsky, M., Small, S. L. & Rauschecker, J. P. Neurobiological roots of language in primate audition: common computational properties. Trends in Cognitive Sciences 19, 142–150 (2015).

30. Kuperberg, G. R. & Jaeger, T. F. What do we mean by prediction in language comprehension? Language, Cognition and Neuroscience 31, 32–59 (2016).

31. Arnal, L. H., Wyart, V. & Giraud, A.-L. Transitions in neural oscillations reflect prediction errors generated in audiovisual speech. Nat Neurosci 14, 797–801 (2011).

32. Blank, H. & Davis, M. H. Prediction Errors but Not Sharpened Signals Simulate Multivoxel fMRI Patterns during Speech Perception. PLoS Biol 14, e1002577 (2016).

33. Kandylaki, K. D. et al. Predicting ‘When’ in Discourse Engages the Human Dorsal Auditory Stream: An fMRI Study Using Naturalistic Stories. Journal of Neuroscience 36, 12180–12191 (2016).

34. Chien, H.-Y. S. & Honey, C. J. Constructing and Forgetting Temporal Context in the Human Cerebral Cortex. Neuron 106, 675–686.e11 (2020).

35. Zadbood, A., Chen, J., Leong, Y. C., Norman, K. A. & Hasson, U. How We Transmit Memories to Other Brains: Constructing Shared Neural Representations Via Communication. Cerebral Cortex 27, 4988–5000 (2017).

36. Baldassano, C. et al. Discovering Event Structure in Continuous Narrative Perception and Memory. Neuron 95, 709–721.e5 (2017).

37. Hamilton, L. S. & Huth, A. G. The revolution will not be controlled: natural stimuli in speech neuroscience. Language, Cognition and Neuroscience 1–10 (2018) doi:10.1080/23273798.2018.1499946.

38. Cohen, J. D. et al. Computational approaches to fMRI analysis. Nat Neurosci 20, 304–313 (2017).

39. Cichy, R. M. & Kaiser, D. Deep Neural Networks as Scientific Models. Trends in Cognitive Sciences 23, 305–317 (2019).

40. Erb, J., Schmitt, L.-M. & Obleser, J. Temporal selectivity declines in the aging human auditory cortex. eLife 9, e55300 (2020).

41. Hochreiter, S. & Schmidhuber, J. Long short-term memory. (1997).

42. Chung, J., Ahn, S. & Bengio, Y. Hierarchical Multiscale Recurrent Neural Networks. arXiv:1609.01704 [cs] (2016).

43. Hale, J. A probabilistic earley parser as a psycholinguistic model, in Proceedings of the North American association of computational linguistics 159–166 (Association for Computational Linguistics, 2001). doi:10.3115/1073336.1073357.

44. Levy, R. Expectation-based syntactic comprehension. Cognition 106, 1126–1177 (2008).

45. Nastase, S. A., Gazzola, V., Hasson, U. & Keysers, C. Measuring shared responses across subjects using intersubject correlation. Social Cognitive and Affective Neuroscience nsz037 (2019) doi:10.1093/scan/nsz037.

46. Glasser, M. F. et al. A multi-modal parcellation of human cerebral cortex. Nature 536, 171–178 (2016).

47. Boldt, R. et al. Listening to an Audio Drama Activates Two Processing Networks, One for All Sounds, Another Exclusively for Speech. PLoS ONE 8, e64489 (2013).

48. Schmälzle, R., Hacker, F. E. K., Honey, C. J. & Hasson, U. Engaged listeners: shared neural processing of powerful political speeches. Social Cognitive and Affective Neuroscience 10, 1137–1143 (2015).

49. Regev, M. et al. Propagation of Information Along the Cortical Hierarchy as a Function of Attention While Reading and Listening to Stories. Cerebral Cortex 29, 4017–4034 (2018).

50. Hasson, U., Yang, E., Vallines, I., Heeger, D. J. & Rubin, N. A Hierarchy of Temporal Receptive Windows in Human Cortex. Journal of Neuroscience 28, 2539–2550 (2008).

51. Zacks, J. M., Speer, N. K., Swallow, K. M., Braver, T. S. & Reynolds, J. R. Event perception: A mind-brain perspective. Psychological Bulletin 133, 273–293 (2007).

52. Radvansky, G. A. Across the Event Horizon. Curr Dir Psychol Sci 21, 269–272 (2012).

53. Zacks, J. M. et al. Human brain activity time-locked to perceptual event boundaries. Nat Neurosci 4, 651–655 (2001).

54. Ditman, T., Holcomb, P. J. & Kuperberg, G. R. Time travel through language: Temporal shifts rapidly decrease information accessibility during reading. Psychonomic Bulletin & Review 15, 750–756 (2008).

55. Whitney, C. et al. Neural correlates of narrative shifts during auditory story comprehension. NeuroImage 47, 360–366 (2009).

56. Richmond, L. L. & Zacks, J. M. Constructing Experience: Event Models from Perception to Action. Trends in Cognitive Sciences 21, 962–980 (2017).

57. Lieder, F., Stephan, K. E., Daunizeau, J., Garrido, M. I. & Friston, K. J. A Neurocomputational Model of the Mismatch Negativity. PLoS Comput Biol 9, e1003288 (2013).

58. Kumar, S., Kaposvari, P. & Vogels, R. Encoding of Predictable and Unpredictable Stimuli by Inferior Temporal Cortical Neurons. Journal of Cognitive Neuroscience 29, 1445–1454 (2017).

59. Todorovic, A. & de Lange, F. P. Repetition Suppression and Expectation Suppression Are Dissociable in Time in Early Auditory Evoked Fields. Journal of Neuroscience 32, 13389–13395 (2012).

60. Brown, C. & Hagoort, P. The Processing Nature of the N400: Evidence from Masked Priming. Journal of Cognitive Neuroscience 5, 34–44 (1993).

61. Hagoort, P., Baggio, G. & Willems, R. M. Semantic unification, in The cognitive neurosciences 819–836 (MIT Press, 2009).

62. Lee, T. S. & Mumford, D. Hierarchical Bayesian inference in the visual cortex. J. Opt. Soc. Am. A 20, 1434 (2003).

63. Kok, P., Jehee, J. F. M. & de Lange, F. P. Less Is More: Expectation Sharpens Representations in the Primary Visual Cortex. Neuron 75, 265–270 (2012).

64. Rao, R. P. N. & Ballard, D. H. Predictive coding in the visual cortex: a functional interpretation of some extra-classical receptive-field effects. Nat Neurosci 2, 79–87 (1999).

65. Bell, A. H., Summerfield, C., Morin, E. L., Malecek, N. J. & Ungerleider, L. G. Encoding of Stimulus Probability in Macaque Inferior Temporal Cortex. Current Biology 26, 2280–2290 (2016).

66. Hickok, G. & Poeppel, D. Dorsal and ventral streams: a framework for understanding aspects of the functional anatomy of language. Cognition 92, 67–99 (2004).

67. Hickok, G. & Poeppel, D. The cortical organization of speech processing. Nature Reviews Neuroscience 8, 393–402 (2007).

68. DeWitt, I. & Rauschecker, J. P. Phoneme and word recognition in the auditory ventral stream. Proceedings of the National Academy of Sciences 109, E505–E514 (2012).

69. Bornkessel-Schlesewsky, I. & Schlesewsky, M. Reconciling time, space and function: A new dorsal-ventral stream model of sentence comprehension. Brain and Language 125, 60–76 (2013).

70. Wilson, S. M., Molnar-Szakacs, I. & Iacoboni, M. Beyond Superior Temporal Cortex: Intersubject Correlations in Narrative Speech Comprehension. Cerebral Cortex 18, 230–242 (2008).

71. Garrido, M. I., Rowe, E. G., Halász, V. & Mattingley, J. B. Bayesian Mapping Reveals That Attention Boosts Neural Responses to Predicted and Unpredicted Stimuli. Cerebral Cortex 28, 1771–1782 (2018).

72. Phillips, H. N. et al. Convergent evidence for hierarchical prediction networks from human electrocorticography and magnetoencephalography. Cortex 82, 192–205 (2016).

73. Meyniel, F. & Dehaene, S. Brain networks for confidence weighting and hierarchical inference during probabilistic learning. Proc Natl Acad Sci USA 114, E3859–E3868 (2017).

74. Cheung, V. K. M., Meyer, L., Friederici, A. D. & Koelsch, S. The right inferior frontal gyrus processes nested non-local dependencies in music. Scientific Reports 8,(2018).

75. Bottini, R. & Doeller, C. F. Knowledge Across Reference Frames: Cognitive Maps and Image Spaces. Trends in Cognitive Sciences 24, 606–619 (2020).

76. Spiers, H. J., Hayman, R. M. A., Jovalekic, A., Marozzi, E. & Jeffery, K. J. Place Field Repetition and Purely Local Remapping in a Multicompartment Environment. Cerebral Cortex 25, 10–25 (2015).

77. Brunec, I. K., Moscovitch, M. & Barense, M. D. Boundaries Shape Cognitive Representations of Spaces and Events. Trends in Cognitive Sciences 22, 637–650 (2018).

78. Alexander, A. S. & Nitz, D. A. Spatially Periodic Activation Patterns of Retrosplenial Cortex Encode Route Sub-spaces and Distance Traveled. Current Biology 27, 1551–1560.e4 (2017).

79. Johnson, A. & Redish, A. D. Neural Ensembles in CA3 Transiently Encode Paths Forward of the Animal at a Decision Point. Journal of Neuroscience 27, 12176–12189 (2007).

80. Stachenfeld, K. L., Botvinick, M. M. & Gershman, S. J. The hippocampus as a predictive map. Nat Neurosci 20, 1643–1653 (2017).

81. Kurby, C. A. & Zacks, J. M. Starting from scratch and building brick by brick in comprehension. Mem Cogn 40, 812–826 (2012).

82. Sainburg, T., Theilman, B., Thielk, M. & Gentner, T. Q. Parallels in the sequential organization of birdsong and human speech. Nat Commun 10, 3636 (2019).

83. Power, J. D., Barnes, K. A., Snyder, A. Z., Schlaggar, B. L. & Petersen, S. E. Spurious but systematic correlations in functional connectivity MRI networks arise from subject motion. NeuroImage 59, 2142–2154 (2012).

84. Rysop, A. U., Schmitt, L.-M., Obleser, J. & Hartwigsen, G. Neural modelling of the semantic predictability gain under challenging listening conditions. Human Brain Mapping 42, 110–127 (2021).

85. McDermott, J. H. & Simoncelli, E. P. Sound Texture Perception via Statistics of the Auditory Periphery: Evidence from Sound Synthesis. Neuron 71, 926–940 (2011).

86. Kisler, T., Reichel, U. & Schiel, F. Multilingual processing of speech via web services. Computer Speech & Language 45, 326–347 (2017).

87. Brysbaert, M. et al. The Word Frequency Effect: A Review of Recent Developments and Implications for the Choice of Frequency Estimates in German. Experimental Psychology 58, 412–424 (2011).

88. van Heuven, W. J. B., Mandera, P., Keuleers, E. & Brysbaert, M. Subtlex-UK: A New and Improved Word Frequency Database for British English. Quarterly Journal of Experimental Psychology 67, 1176–1190 (2014).

89. Brainard, D. H. The Psychophysics Toolbox. Spatial Vis 10, 433–436 (1997).

90. Heinzerling, B. & Strube, M. BPEmb: Tokenization-free Pre-trained Subword Embeddings in 275 Languages. arXiv:1710.02187[cs] (2017).

91. Kádár, Á., Côté, M.-A., Chrupała, G. & Alishahi, A. Revisiting the Hierarchical Multiscale LSTM. arXiv:1807.03595[cs] (2018).

92. Kingma, D. P. & Ba, J. Adam: A Method for Stochastic Optimization. arXiv:1412.6980 [cs] (2017).

93. Barbaresi, A. A corpus of German political speeches from the 21st century. 11th Language Resources and Evaluation Conference 792–797 (2018).

94. Abadi, M. et al. TensorFlow: Large-Scale Machine Learning on Heterogeneous Distributed Systems. (2015).

95. Broderick, M. P., Anderson, A. J., Di Liberto, G. M., Crosse, M. J. & Lalor, E. C. Electrophysiological Correlates of Semantic Dissimilarity Reflect the Comprehension of Natural, Narrative Speech. Current Biology 28, 803–809.e3 (2018).

96. Chi, T., Ru, P. & Shamma, S. A. Multiresolution spectrotemporal analysis of complex sounds. The Journal of the Acoustical Society of America 118, 887–906 (2005).

97. Kumar, M. et al. BrainlAK tutorials: User-friendly learning materials for advanced fMRI analysis. PLoS Comput Biol 16, e1007549 (2020).

98. Benjamini, Y. & Hochberg, Y. Controlling the False Discovery Rate: A Practical and Powerful Approach to Multiple Testing. Journal of the Royal Statistical Society, Series B (Methodological) 57, 289–300 (1995).

99. Santoro, R. et al. Reconstructing the spectrotemporal modulations of real-life sounds from fMRI response patterns. Proc Natl Acad Sci USA 114, 4799–4804 (2017).

100. Golub, G. H., Heath, M. & Wahba, G. Generalized Cross-Validation as a Method for Choosing a Good Ridge Parameter. Technometrics 21, 215–223 (1979).

101. Musall, S., Kaufman, M.T., Juavinett, A. L., Gluf, S. & Churchland, A. K. Single-trial neural dynamics are dominated by richly varied movements. Nat Neurosci 22, 1677–1686 (2019).

102. Pedregosa, E. et al. Scikit-learn: Machine Learning in Python. Journal of Machine Learning Research 12, 2825–2830 (2011).

103. Maris, E. & Oostenveld, R. Nonparametric statistical testing of EEG- and MEG-data. Journal of Neuroscience Methods 164, 177–190 (2007).

104. Friston, K. J. et al. Psychophysiological and Modulatory Interactions in Neuroimaging. NeuroImage 6, 218–229 (1997).

105. Pennington, J., Socher, R. & Manning, C. Glove: Global Vectors for Word Representation, in Proceedings of the 2014 Conference on Empirical Methods in Natural Language Processing (EMNLP) 1532–1543 (Association for Computational Linguistics, 2014). doi:10.3115/v1/D14-1162.

106. Ba, J. L., Kiros, J. R. & Hinton, G. E. Layer Normalization. arXiv:1607.06450[cs, stat] (2016).

107. Penny, W., Friston, K., Ashburner, J., Kiebel, S. & Nichols, T. Statistical Parametric Mapping: The Analysis of Functional Brain Images. (Academic Press, 2006).

108. Esteban, O. et al. fMRIPrep: a robust preprocessing pipeline for functional MRI. Nature Methods 16, 111–116 (2019).

109. Gorgolewski, K. et al. Nipype: A Flexible, Lightweight and Extensible Neuroimaging Data Processing Framework in Python. Frontiers in Neuroinformatics 5, (2011).

110. Abraham, A. et al. Machine learning for neuroimaging with scikit-learn. Frontiers in Neuroinformatics 8,(2014).

111. Tustison, N. J. et al. N4ITK: Improved N3 Bias Correction. IEEE Transactions on Medical Imaging 29, 1310–1320 (2010).

112. Dale, A. M., Fischl, B. & Sereno, M. I. Cortical Surface-Based Analysis. NeuroImage 9, 179–194 (1999).

113. Fonov, V., Evans, A., McKinstry, R., Almli, C. & Collins, D. Unbiased nonlinear average age-appropriate brain templates from birth to adulthood. NeuroImage 47, S102 (2009).

114. Avants, B., Epstein, C., Grossman, M. & Gee, J. Symmetric diffeomorphic image registration with cross-correlation: Evaluating automated labeling of elderly and neurodegenerative brain. Medical Image Analysis 12, 26–41 (2008).

115. Klein, A. et al. Mindboggling morphometry of human brains. PLoS Comput Biol 13, e1005350 (2017).

116. Zhang, Y., Brady, M. & Smith, S. Segmentation of brain MR images through a hidden Markov random field model and the expectation-maximization algorithm. IEEE Trans. Med. Imaging 20, 45–57 (2001).

117. Jenkinson, M., Bannister, P., Brady, M. & Smith, S. Improved Optimization for the Robust and Accurate Linear Registration and Motion Correction of Brain Images. NeuroImage 17, 825–841 (2002).

118. Cox, R. W. AFNI: Software for Analysis and Visualization of Functional Magnetic Resonance Neuroimages. Computers and Biomedical Research 29, 162–173 (1996).

119. Greve, D. N. & Fischl, B. Accurate and robust brain image alignment using boundary-based registration. NeuroImage 48, 63–72 (2009).

120. Pruim, R. H. R. et al. ICA-AROMA: A robust ICA-based strategy for removing motion artifacts from fMRI data. NeuroImage 112, 267–277 (2015).

121. Power, J. D. et al. Methods to detect, characterize, and remove motion artifact in resting state fMRI. NeuroImage 84, 320–341 (2014).

122. Lindquist, M. A., Geuter, S., Wager, T. D. & Caffo, B. S. Modular preprocessing pipelines can reintroduce artifacts into fMRI data. Human Brain Mapping 40, 2358–2376 (2019).

123. Guntupalli, J. S. et al. A Model of Representational Spaces in Human Cortex. Cereb. Cortex 26, 2919–2934 (2016).

124. Hanke, M. et al. PyMVPA: a Python Toolbox for Multivariate Pattern Analysis of fMRI Data. Neuroinform 7, 37–53 (2009).

