## Supplementary Materials for "Predicting speech from a cortical hierarchy of event-based timescales"

### Supplementary Text

#### Supplementary Text 1: Mapping words to pre-trained BPEmb subword vectors

In short, the BPEmb vocabulary is based on a text corpus, which was segmented into its subwords using byte-pair encoding. That is, smaller subword units (e.g., letters) most frequently co-occurring in the corpus were iteratively merged into larger units (e.g., syllables) and added to the vocabulary until the predefined maximum of merge operations was reached (i.e., vocabulary size). The corresponding embeddings were trained with the GloVe algorithm[105]. Importantly, the length of subwords ranged from single letters to complete words. For example, the inflected verb “fischte” [“fished”; 3rd person singular, simple past, active voice, indicative of “to fish”] consists of one subword embedding representing its word stem “fisch” and another embedding representing its suffix “te”, whereas the more frequent word “Wasser” [“water”] is represented by only one embedding.

#### Supplementary Text 2: Architecture of language models

In a simple recurrent neural network (RNN), the hidden state $h_{p-1}$ stores all relevant context and is sequentially passed to the next cell where it is updated with information from word $w_p$. More specifically, the recurrent input and the bottom-up input are combined into the cell input vector $g_p$:

$$g_p = tanh\left(W_g w_p + U_g h_{p-1} + b_g\right),$$

where *tanh* is the activation function, $W_g \in R^{n\times e}$ and $U_g \in R^{n\times n}$ are trainable weight matrices, *n* is the number of neurons (or units), $b \in R^{n\times 1}$ is a bias term, $w_p \in R^{e\times 1}$ and $h_{p-1} \in R^{n\times 1}$.

In the continuously updating LSTM, the cell state $c_p$ acts as long-term memory, whereas the hidden state $h_p$ incorporates information relevant to the cell output (i.e., the prediction of the next word). The integration of new information and the information flow between the two memory systems is controlled by three gating mechanisms. First, the cell state is updated. The forget gate $f_p$ determines which information from the previous cell state $c_{p-1}$ has become irrelevant and should be removed by:

$$f_p = \sigma\left(W_f w_p + U_f h_{p-1} + b_f\right),$$

where $\sigma$ is the sigmoid activation function. The input gate $i_p$ determines which information from candidate state $g_p$ should be added to the cell state by:

$$i_p = \sigma\left(W_i w_p + U_i h_{p-1} + b_i\right).$$

The new cell state $c_p$ is created by:

$$c_p = f_p \odot c_{p-1} + i_p \odot g_p,$$

where $c_{p-1} \in R^{n \times 1}$. Second, the hidden state is updated. The output gate $o_p$ determines which information from long-term memory $c_p$ might become relevant shortly and should be added to the hidden state by:

$$o_p = \sigma(W_o w_p + U_o h_{p-1} + b_o).$$

The new hidden state $h_p$ is created by:

$$h_p = o_p \odot tanh(c_p).$$

The sparsely-updating HM-LSTM employs a revised updating rule where information from the lower layer is only fed forward at the end of a timescale (i.e., a sequence of words closely related to each other). To this aim, $z_p^l$ is introduced which marks the end of a timescale:

$$\tilde{z}_p^l = hard\ sigm(z_p^{l-p} W_z h_p^{l-1} + U_z h_{p-1}^l + b_z^l),$$

$$z_p^l = \begin{cases} 1 & \text{if } \tilde{z}_p^l > 0.5 \\ 0 & \text{otherwise,} \end{cases}$$

where *hard sigm* is the hard sigmoid activation function. If $z_p^{l-1} = 1$, the hidden state $h_p^l$ and cell state $c_p^l$ are computed like in a vanilla LSTM cell ("update mechanism"). Otherwise, the hidden state $h_p^l$ and cell state $c_p^l$ are simply the copy of $h_{p-1}^l$ and $c_{p-1}^l$ ("copy mechanism"), respectively.

**Supplementary Text 3: Prediction of the next word**

LSTM and HM-LSTM cells form the representations of semantic information relevant to speech prediction, whereas the actual prediction of the next word takes place in the output module. Here, hidden states at word position *p* are combined across the different layers of the language model by:

$$h_p^r = LReLU\left(\sum_{l=1}^{L} W_r^l h_p^l\right),$$

where *LReLU* is the leaky rectified linear unit activation function and *L* is the number of layers. The combined hidden state $h_p^r$ is mapped to a fully connected dense layer of as many neurons as there are words in the vocabulary and squashed to values in the interval [0,1], which sum to 1:

$$d_p = softmax(W_d h_p^r + b_d),$$

where *softmax* is the squashing function, $W_d \in R^{v \times n}$ and $b_d \in R^{v \times 1}$. Each neuron in vector $d_p$ indexes one particular word in vocabulary *v* and denotes its probability of being the next word. Finally, the word referring to the highest probability in the distribution is chosen by:

$$s_p = argmax(d_p),$$

where $s_p$ is the predicted next word in a story.

#### Supplementary Text 4: Training language models

All other architectural choices were based on results from systematic ablation tests on HM-LSTM cells[91]. Accordingly, the optimizer's initial learning rate of 0.001 was reduced by a factor of 0.02 when performance on the validation set did not improve over one epoch. Gradients were clipped at a value of 1. We applied layer normalization to all inputs after multiplication with their respective weight matrices[106]. Further, we added an $l^2$-norm penalty term for weight size to the loss function ($\lambda$ = 0.0005). The dense layer in the output module was excluded from normalization and regularization.

#### Supplementary Text 5: Convolving features with the hemodynamic response function

For features of predictiveness and linguistics, we modelled (higher frequency, randomly spaced) information on the word level as hemodynamic responses sampled at the (lower frequency, equally spaced) TR of fMRI data. This was achieved by creating feature vectors of zeros corresponding to the length of a functional run with a sampling frequency of 1,000 Hz, which allowed word onsets and offsets to be represented with high temporal precision. For each word in a run, a boxcar function, which was scaled to the feature's value at that particular word, was placed on all samples falling in between word onset and offset. The resulting vector including feature values for all words in a run was convolved with SPM's canonical hemodynamic response function (HRF)[107] and downsampled to the TR. Acoustic features, on the other hand, were already sampled to the TR and therefore directly convolved with the HRF.

#### Supplementary Text 6: Preprocessing structural and functional MRI

**Structural MRI data preprocessing.** MRI data were preprocessed with fMRIPrep 1.2.4[108], which is based on Nipype 1.1.6[109] and employs Nilearn 0.5.0[110] in many internal operations. For each participant, the T1w image was corrected for intensity non-uniformity using N4BiasFieldCorrection (ANTs 2.1.0)[111] and then skull-stripped using the OASIS template in antsBrainExtraction.sh (ANTs 2.2.0). Individual brain surfaces were reconstructed from T1w and T2w reference images using recon-all (FreeSurfer 6.0.1)[112]. The T1w reference image was spatially normalized to the MNI152NLin2009cAsym template[113] through nonlinear registration with antsRegistration (ANTs 2.2.0)[114].

A brain mask was created by reconciling ANTs-derived and FreeSurfer-derived segmentations of the cortical grey matter according to a customized variation of the implementation in Mindboggle[115]. Brain tissue segmentation of cerebrospinal fluid, white matter and grey matter was performed on the T1w reference image using FAST (FSL 5.0.9)[116].

**Functional MRI data preprocessing.** For each functional run, BOLD time series were motion corrected using mcflirt (FSL 5.0.9)[117] and slice time corrected using 3dTshift (AFNI 20160207)[118]. After unwarping BOLD images based on the susceptibility distortion estimated from field maps, the BOLD reference image was aligned to the native T1w reference image using boundary-based registration with six degrees of freedom[119] as implemented in bbregister (FreeSurfer). BOLD images were resampled to standard space. To correct for head motion, non-aggressive Automatic Removal of Motion Artifacts using Independent Component Analysis (ICA-AROMA)[120] was performed on the resampled and smoothed (6 mm FWHM Gaussian kernel) BOLD images. On average, 50.54 % of the maximal 200 components per functional run ($Ra = 32.93–71.78$, $SD = 9.21$) were classified as motion-related artefacts. Additional confounding noise time series like the average signal within cerebrospinal fluid and white matter as well as framewise displacement were calculated in Nipype following the definitions by Power and colleagues[121]. A Discrete Cosine Transform (DCT) basis set of six functions with a cut-off at 0.008 Hz was generated for temporal high-pass filtering.

After running fMRIPrep, we regressed out high-pass filters as well as cerebrospinal fluid and white matter signals from the BOLD time series using 3dTproject (AFNI 19.2.24). To avoid reintroducing previously removed artefacts into the functional data[122], we projected the ICA-AROMA artefact components onto the additional nuisance covariates and used the residuals as predictors orthogonal to prior predictors. The denoised BOLD images were resampled to the fsaverage5 template in surface space by averaging across the cortical ribbon in 5 equally spaced steps at each vertex using trilinear interpolation. All resamplings can be performed with a single interpolation step: volumetric resamplings were performed using antsApplyTransforms (ANTs 2.1.0) with Lanczos interpolation; surface resamplings were performed using mri_vol2surf (FreeSurfer). In each functional run, the first 10 baseline volumes as well as the last volume were removed from time series and all further analysis were carried out on *z*-scored single-vertex BOLD time series.

**Functional alignment to a common space.** To account for small spatial variations in intersubject response tuning, functional time series were projected into a common space using searchlight hyperalignment across the whole cortex as described by Guntupalli and colleagues[123]. Here, we centred a sphere (or searchlight) with a radius of 20 mm on each vertex and determined the optimal rotation of response vectors (or functional time series) within each searchlight in three iterations using Procrustes transformation. An intermediate common space was initialized by rotating one participant's response vectors to best match the responses of a randomly chosen reference participant. All other participants were successively brought into alignment, with the average of all previously rotated response vectors as a reference. In a second iteration, all original response vectors were aligned to the intermediate common space and the average of resulting rotated response vectors became the final common space. In the third iteration, hyperalignment parameters mapping single participant's original response vectors to the final common space were calculated. Parameters corresponding to vertices of overlapping searchlights were averaged. We ran hyperalignment on four independent data splits (i.e., pairing up every fourth of eight functional runs) and averaged transformation matrices across data splits to derive final parameters for each participant. Hyperalignment was performed in PyMVPA (2.6.6)[124].

### Supplementary figures

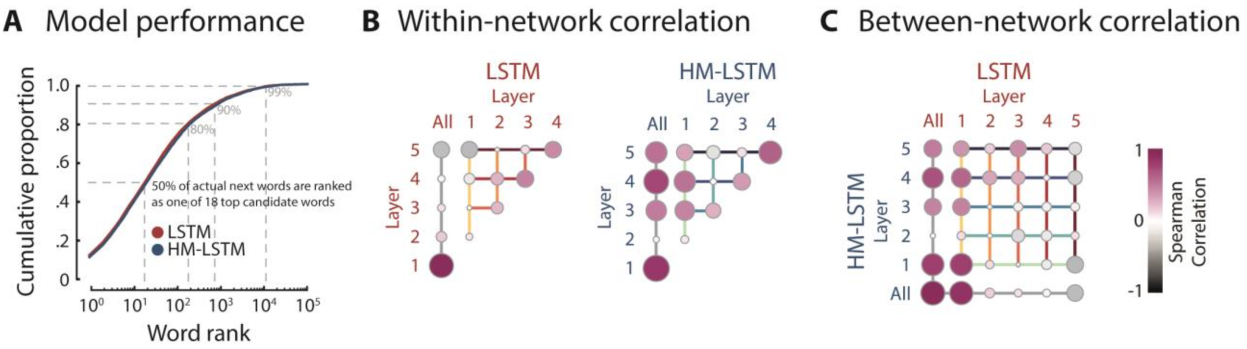


**Supplementary Figure 1. Evaluating language models. (A)** For each language model (LSTM: red, HM-LSTM: blue), we extracted the rank of the next word from the probability distribution of all candidate words. Language models ranked more than 50 % of words in the text as one of 18 top candidates words (out of more than 90,000 words). **(B)** Spearman correlations of word surprisal between single layers of language models, separately for LSTM (left) and HM-LSTM (right). In addition, correlations of single layers with full models are shown; color and size of circles scale to correlation coefficients. **(C)** Spearman correlations of word surprisal between language models.

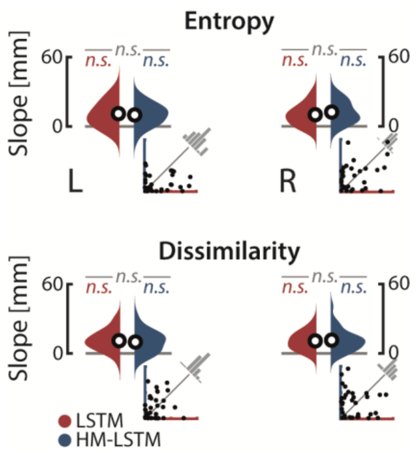


**Supplementary Figure 2. Encoding the timescales of entropy and dissimilarity.** Along the dorsal stream, linear functions were fit to peak coordinates of entropy (top) and dissimilarity (bottom) across timescales. Resulting slope parameters were compared to null distributions drawn from scrambled coordinates and between language models (LSTM: red, HM-LSTM: blue), separately for the left (left column) and right hemisphere (right column). Black circles represent grand-average slope parameters; insets depict coefficients of determination for linear fits of single participants. *n.s.*: not significant.
